# Crystal structures and fragment screening of SARS-CoV-2 NSP14 reveal details of exoribonuclease activation and mRNA capping and provide starting points for antiviral drug development

**DOI:** 10.1101/2022.03.11.483836

**Authors:** Nergis Imprachim, Yuliana Yosaatmadja, Joseph A Newman

## Abstract

The SARS-CoV-2 non-structural protein 14 (NSP14) is a dual function enzyme containing an N-terminal exonuclease domain (ExoN) and C-terminal Guanine-N7-methyltransferase (N7-MTase) domain. Both enzymatic activities appear to be essential for the viral life cycle and thus may be targeted for anti-viral therapeutics. NSP14 forms a stable complex with the SARS-CoV-2 zinc binding protein NSP10, and this interaction greatly enhances the nuclease but not the methyltransferase activity. In this study, we have determined the crystal structure of SARS-CoV-2 NSP14 in the absence of NSP10 to 1.7 Å resolution. Comparisons of this structure with the structure of NSP14/NSP10 complexes solved to date reveal significant conformational changes that occur within the NSP14 ExoN domain upon binding of NSP10, including significant movements and helix to coil transitions that facilitate the formation of the ExoN active site and provide an explanation of the stimulation of nuclease activity by NSP10. Conformational changes are also seen in the MTase active site within a SAM/SAH interacting loop that plays a key role in viral mRNA capping. We have also determined the structure of NSP14 in complex with cap analogue ^7Me^GpppG, offering new insights into MTase enzymatic activity. We have used our high resolution crystals to perform X-ray fragment screening of NSP14, revealing 72 hits bound to potential sites of inhibition of the ExoN and MTase domains. These structures serve as excellent starting point tools for structure guided development and optimization of NSP14 inhibitors that may be used to treat COVID-19 and potentially other future viral threats.

## Introduction

The global COVID-19 pandemic is caused by the SARS-CoV-2 virus and has infected over 200 million and caused over 4 million fatalities worldwide at the point of writing. Despite the successes with vaccines, there are currently a lack of effective drugs to treat people infected with SARS-CoV-2, and identification of such agents is a global priority. SARS-CoV-2 has a relatively large (∼30 Kb) positive-sense single stranded RNA genome that consists of 12 functional open reading frames (ORF), encoding four structural proteins, six accessory proteins, and 16 non-structural proteins NSP1-16^1,2^. The non-structural proteins (NSPs) consist of the machinery for viral replication including RNA dependent RNA polymerase, primase, proteases, helicase, nucleases and various enzymes involved in RNA capping.

The SARS-CoV-2 NSP14 is a bifunctional protein containing an N-terminal DEDDh-type exoribonuclease (ExoN) domain and a C-terminal SAM-dependent guanine-N7 methyltransferase (N7-MTase) domain^3^. The ExoN domain is thought to play a role in RNA proofreading and maintaining the integrity of the viral genome by removal of mismatched nucleotides in a metal ion dependent manner^4,5^. This exonuclease activity is greatly enhanced by the interaction of NSP14 with its activator protein NSP10. NSP10 is a small 139 amino acid with a mixed α/β fold that does not resemble any know prokaryotic or eukaryotic homologues and coordinates two structural zinc ions via C3H1 and C4 zinc binding motifs^6,7^. Structures of SARS CoV NSP14/NSP10 complexes have been solved by crystallography^5^, and more recently the SARS-CoV-2 NSP14-ExoN domain of in complex with NSP10^8^, as well as the NSP14/NSP10 in complex with a mismatch containing RNA by cryo-EM^9^. These structures show an extensive predominantly hydrophobic interface (in excess of 2000 Å^2^) with the N-terminal region of NSP14 packing against two faces of NSP10. The metal active site of NSP14 lies close to the NSP10 interacting regions although a structure of NSP14 in the absence of NSP10 is not known and thus the molecular basis for the nuclease activity stimulation is not fully understood.

The NSP14 N7-MTase domain is located in the C-terminus of NSP14 (residues 288-527) and contributes an important methylation step in the formation of the viral 5’ RNA cap, which is essential for hijacking host translational machinery and suppressing innate immune responses. The viral 5’ RNA cap is formed by the sequential action of multiple enzymes that collectively assemble into a multi protein Replication and Transcription Complex (RTC). The helicase NSP13 is the RNA 5’ triphosphatase that cleaves the terminal triphosphate to a diphosphate^10^. The SARS-CoV-2 RNA dependent RNA polymerase NSP12 NiRAN domain was recently identified as the guanylyltransferase that enzyme that transfers GMP from GTP to the diphosphate to form a GpppN structure^11^. NSP14 then methylates the guanine N7 of the GpppN structure in a S-adenosyl methionine (SAM) dependent manner to form cap-0 structure (^7Me^GpppN)^12^. The product of this reaction is further methylated at the ribose O2’ position by NSP16 to form the final cap-1 structure (^7Me^GpppN_2’OMe_)^13^, which provides viral RNA stability and host immune evasion. Studies suggest that this 2′-O-MTase activity of NSP16 is stimulated by NSP10^14^. NSP16 forms a complex with NSP10 via distinct yet overlapping interface. The apparent stoichiometric discrepancy is relieved by the fact that NSP10 is present on both open reading frames ORF1a and ORF1ab unlike NSP14 and NSP16. Recent cryo-EM structures have shown how the dual enzymatic activities of NSP14 are coupled to the rest of the RTC machinery via a physical protein interaction between the NSP14 ExoN domain and NSP9 and NSP12 NiRAN domain within the RTC^14^. This interaction suggests the existence of a co-transcriptional capping complex (NSP12/NSP9/NSP14/NSP10) and the authors suggest that the ExoN domain may function in an *in trans* backtracking mechanism for proofreading.

In this study, we have determined the 1.7 Å crystal structure of SARS-CoV-2 NSP14 alone in the absence of its partner NSP10. Unexpectedly significant regions of the interaction interface retain some degree of folded structure. The predominantly hydrophobic interface, however, forms a collapsed self-association with significant rearrangement of the loops and residues that form the ExoN active site. The MTase domain also shows conformational differences when compared with previously solved NSP14/NSP10 structures. The MTase SAM binding loop occupies an alternative conformation that may be optimal for binding the ^7Me^GpppN product. NSP14 is one of the most conserved targets in the Coronavirus genome and thus is an excellent target for broad spectrum antivirals. We have used our crystals to perform a crystallographic fragment screen of NSP14 and find multiple fragments bound to sites of inhibition including the MTase and ExoN active sites and sites that may block the potential of NSP14 to interact with its partner NSP10. These fragments are important starting points for structure guided development or optimization of NSP14 inhibitors that may be useful as antiviral therapeutics with the potential for broad spectrum of action.

## Results and Discussion

### Crystal structure of NSP14 at 1.7 Å resolution

Initial attempts to crystallize the SARS-CoV-2 NSP14/NSP10 complex, either expressed in *E*.*coli* as a bicistronic construct, or reconstituted from components *in vitro*, failed to produce crystals of diffraction quality. We explored N-terminal truncations of NSP14 as a way to increase stability of NSP14. We found a construct omitting the first 7 amino acids (residues 8-527) gave significantly improved purification yields. Crystallization of this NSP14 construct alone in a condition containing 1.26 M sodium phosphate monobasic and 0.14 M potassium phosphate dibasic produced rod shaped crystals that diffracted to 1.7 Å with moderately anisotropic diffraction. We solved the structure by molecular replacement using NSP14 from SARS-CoV NSP14/NSP10 complex^5^ as the starting model (PDB entry 5C8U). Overall, the electron density map is of high quality (Figure S1) and the model includes 3 Zinc ions and 2 Phosphate ions. The NSP14 model is complete with the exception of 17 residues in the N-terminus, 3 in the C-terminus and 3 loops (amino acids 95-102, 122-151 and 456-463), which are not visible in the electron density presumably due to disorder. The model is of high quality with good stereochemistry and has been refined to a final R=0.197 R_free_=0.220. A summary of the data collection and refinement statistics are shown on Table I.

**Table 1.** Data collection and refinement statistics.

| Table 1 Data collection and refinement statistics |  |  |
| --- | --- | --- |
|  | NSP14 | NSP14-7MeGpppG |
| Space group | P212121 | P212121 |
| Cell dimensions, *a,b,c* (Å) | 69.2, 68.1, 138.5 | 67.3, 67.3, 138.6 |
| Angles $\alpha,\beta,\gamma$ (°) | 90, 90, 90 | 90, 90, 90 |
| Wavelength (Å) | 0.9179 | 0.97625 |
| Resolution (Å) | 69.2-1.66 (1.850-1.66) | 69.3 – 2.53 (2.73 – 2.53) |
| R<sub>merge</sub> | 0.125 (1.075) | 0.585 (1.990) |
| I/σI | 9.2 (1.6) | 6.0 (1.6) |
| CC1/2 | 0.997 (0.716) | 0.986 (0.747) |
| Completeness (spherical) % | 49.0 (8.7) | 55.5 (13.6) |
| Completeness (ellipsoidal) % | 89.27 (65.6) | 91.7 (65.9) |
| Multiplicity | 5.5 (6.2) | 26.5 (27.6) |
| No. Unique reflections | 37654 (1884) | 12041 (602) |
| Refinement statistics |  |  |
| Resolution | 69.2 - 1.66 | 69.3 – 2.53 |
| R<sub>work</sub>/R<sub>free</sub> (%) | 19.7/22.0 | 21.2/24.1 |
| No. atoms | 4116 | 3782 |
| Protein | 3672 | 3672 |
| Solvent | 431 | 45 |
| Ligand/ion | 5 | 65 |
| Average B factors (Å<sup>2</sup>) |  |  |
| All atoms | 28.0 | 26 |
| Protein | 27.3 | 25 |
| Solvent | 34.1 | 19 |
| Ligand/ion | 44.0 | 60 |
| Wilson B | 22 | 25 |
| R.M.S. deviations |  |  |
| Bond lengths (Å) | 0.003 | 0.003 |
| Bond angles (°) | 0.551 | 0.762 |
| Ramachandran plot |  |  |
| Favoured (%) | 96.2 | 95.6 |
| Allowed (%) | 3.34 | 3.7 |
| PDB ID | 7QGI | 7QIF |

### Structural comparison of NSP14 alone with NSP14/NSP10 complex

Comparison of our NSP14 structure with previously solved NSP14/NSP10 structures from SARS-CoV-2 or coronavirus species reveals a generally good agreement for the overall structure (overall RMSD values in the range of 1.5 Å) with distinct conformational differences observed for both the ExoN and MTase domains (Figure 1A). The most prominent difference occurs in the fold of the ExoN domain with the regions at the NSP14 N-terminus that interact with NSP10. In the NSP14/NSP10 complex structures, this region forms a small 3 stranded antiparallel β-sheet (strands β1↑β5↓β6↑) which makes extensive contacts to the first (C3H1) zinc ion binding site on NSP10 (Figure 1B &1C). A long region of largely extended coil structure preceding β1 that is interspersed with a small β-hairpin also makes extensive contacts to NSP10 in the region of the NSP10 β-subdomain. This region is also contacted by NSP16 in the NSP16/NSP10 heterodimeric complex^15^. The overall interface covers around 2000 Å^2^ of contact area includes 23 hydrogen bonds, 1 salt bridge and several clusters of buried hydrophobic residues. This binding interface has previously been described as similar to a NSP14 “hand” over NSP10 “fist”^16^ (Figure 1D).

**Figure 1.**
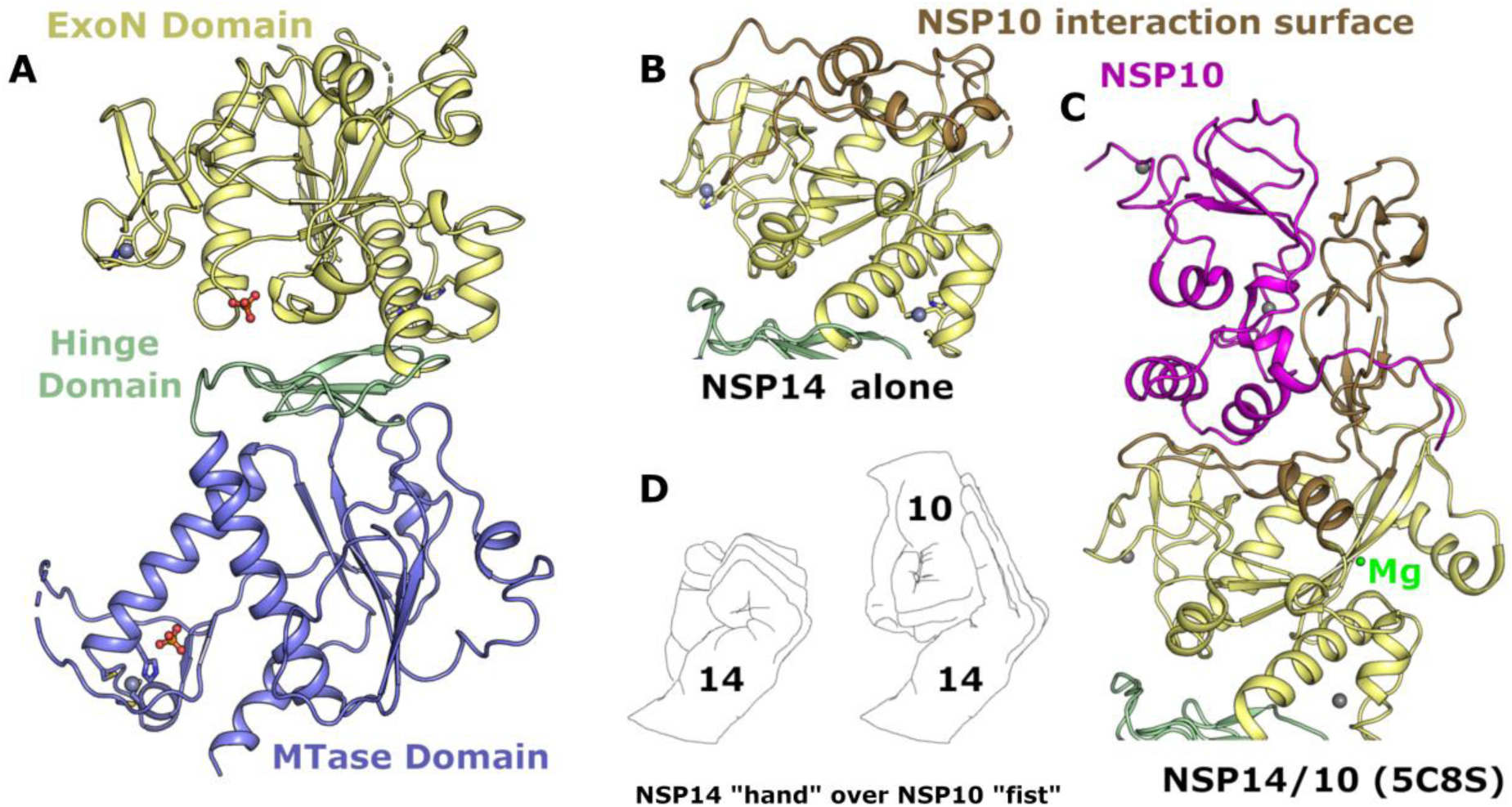
Structure of NSP14 in the absence of NSP10. (**A**) Overall NSP14 structure with the ExoN domain colored yellow, hinge domain in green and MTase domain blue, zinc ions are shown as grey spheres. (**B**) Close up view of the ExoN domain with regions that were observed to shift conformation highlighted in “sand” color. (**C**) Structure of the ExoN domain of SARS-CoV NSP14 in complex with NSP10 (shown in pink) viewed from the same orientation as for panel B. (**D**) The interaction between NSP14 and NSP10 has been described as “hand over fist”, by the same logic the “fingers” of NSP14 collapse back toward the core resembling a closed “fist”.

Somewhat surprisingly, given the intimate association of the N-terminal NSP14 region with NSP10, much this region is still ordered in the absence of NSP10 (103 residues observed out of a possible 153). However, the fold of this region is distinctly different and bears almost no relation to the complex structure (Figure 1B,1C). In particular the first 60 residues forms a more compact arrangement with a higher helical content, and the first β-sheet that associates with the first NSP10 zinc binding site is not formed. Several sequences are observed to transition between secondary structure elements, with residues 36-41 transitioning from coil to helix, residues 54-58 transitioning from β-strand to helix, and the helix 146-152 (second α-helix of the NSP14/10 complex) now partially forms a β-strand and associates with the main β-sheet in NSP14 to form an additional 6^th^ strand (mixed parallel/antiparallel) (Figure S2). This more compact helical N-terminal region associates closely with the remainder of the ExoN domain in a collapsed arrangement such that much of the interface area in the NSP14/10 complex is not solvent exposed but now buried within the new interface between the N-terminal region of NSP14 and the rest of the ExoN domain (Figure 1B). A long stretch of 27 residues spanning amino acids 123-150 are disordered in our NSP14 structure and given the fact that the neighboring residues lie on different ends of the main central ExoN β-sheet (required to span a gap of around 25 Å) may be assumed to adopt a relatively extended coil like fold rather than forming the remaining part of the small N-terminal β-sheet as for the NSP14/10 complex structure.These dramatic conformational changes and collapsing of the N-terminal NSP10 interacting region onto the ExoN core are consistent with predictions made from SAXS analysis of SARS-CoV NSP14 upon binding NSP10^16^. The potential energetic cost of the rearrangements may also explain why the NSP14 - NSP10 interaction appears to have only a moderate affinity (K_d_ value of around 1 µM)^17^ despite the extensive nature of the interface.

### NSP10 complex formation is required to form the ExoN active site

The exonuclease active site is formed in cleft on top of the central ExoN domain β-sheet close to the junction with the N-terminal NSP10 interacting region (Figure 2A & 2B). Four conserved acidic residues D90, E92, E191 and D273 form the metal binding active site core, whilst the fifth catalytic residue H268 has been proposed in related proteins to function as a general base that deprotonates a catalytic water for nucleophilic attack^18^ (Figure 2C). The metal ion binding site has been suggested to bind to two metal ions based on similarities with other enzymes from the family, including exonuclease domains of bacterial polymerases^19^. Current structures in the absence of nucleic acid substrates generally show a single metal ion (typically Mg^2+^) bound in an octahedral environment coordinated with D90, E92, D273 and three water molecules^8,20^. Recent cryo-EM structures of NSP14/NSP10 in complex with RNA^9^ include a second metal ion binding site adjacent to the first coordinated by E191 and D90 and additional two oxygen atoms on the RNA phosphate substrate, suggesting this metal ion may bind in conjunction with RNA (Figure 2C). This study also established the wider NSP14 RNA binding site which contains contributions from the extreme N-terminus of both NSP14 and NSP10, the β2 – β3 loop, the β6 – α2 loop and the two loops preceding and following α5 (Figure 2A). In the absence of NSP10 binding, no metal ion was observed to bind. Indeed, the loop between β7 and α4 is in a different conformation where H188 occupies part of the metal binding pocket and likely prevents the binding of the second metal (Figure 2D). The movement of this loop appears to be influenced by the burying of N-terminal hydrophobic residues F33 and L38 into a hydrophobic environment immediately adjacent to the active site (Figure 2D). Several other regions of the collapsed N-terminal region are in positions where they would form steric clashes with potential substrates (Figure 2B), suggesting that a different mode of RNA binding may be necessary. In the NSP14/NSP10 RNA complex structures, the duplex RNA contains a 5’ overhang with a single separated base pair that has been flipped out by the actions of N104 and H95^9^. Both of these residues are no longer part of the extended active site of NSP14 in the absence of NSP10, and the active site is generally more open, raising the possibility of accommodating fully duplex substrates. The requirement for base separation for access to the NSP14/NSP10 active site has been suggested to confer a preference for mismatched nucleotides^4^ although there is some debate about the existence of such a preference^21^. An intriguing possibility is that NSP14 alone may differ from the NSP14/10 complex in this regard.

**Figure 2.**
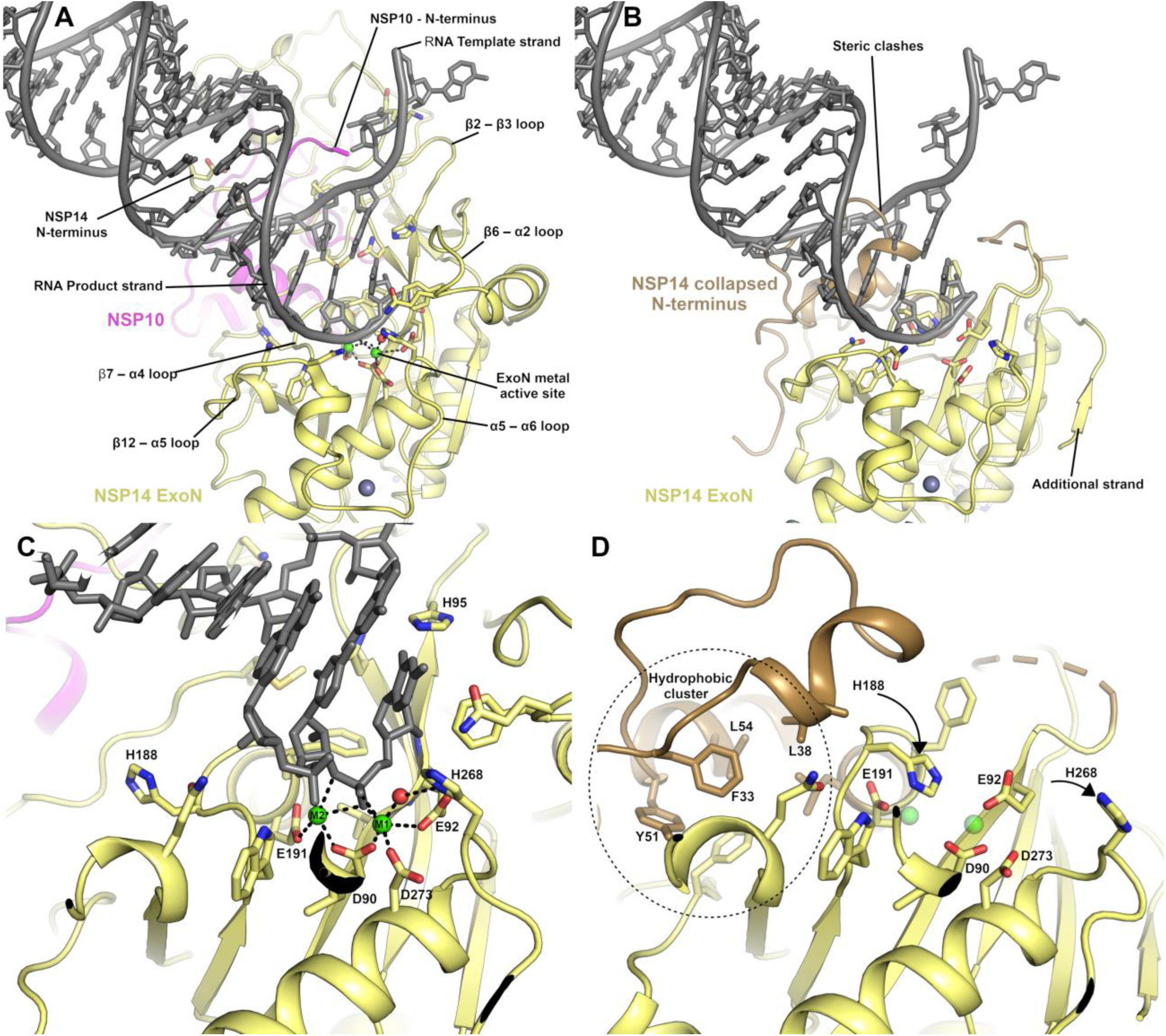
The NSP14 ExoN active site is formed upon binding to NSP10. (**A**) View of the ExoN domain of the NSP14 NSP10 complex bound to a mismatch containing double stranded RNA molecule as recently solved by cryo EM^9^. Key residues in the catalytic center are shown in the stick format and the secondary structure elements that form the wider active site are labelled. (**B**) View of the NSP14 ExoN domain in the absence of NSP10, the mismatch containing RNA is shown for reference and forms significant steric clashes with residues that constitute the collapsed N-terminus (shown in sand color). (**C**) Zoomed in view of the active site catalytic center of the NSP14 NSP10 RNA complex with metal ion coordinating residues labelled. (**D**) NSP14 in the absence of NSP10 viewed from the same orientation. A hydrophobic cluster adjacent to the active site (shown in Sand color) flips H188 into a position where it likely disrupts binding of M2 (shown for reference as semitransparent green spheres).

### The SARS-CoV-2 MTase domain and structure with CAP analogue

The MTase domain of NSP14 is located in the C-terminal half and consists of a central 5 stranded predominantly parallel β-sheet that is capped at its C-terminal end by and additional small 3 stranded antiparallel β-sheet (inserted between the 4^th^ and 5^th^ strands of the main β-sheet) that has been described as a “hinge” domain^16^, as it separates the MTase and ExoN domains (Figure 1A). The MTase fold is distinct from traditional Rossman type α-β-α folds that feature in other known SAM dependent RNA methyltransferases^22^. The MTase active site is formed in a deep pocket between the β-sheets of the MTase and hinge domain with contributions from the long helix (α7) that precedes the first strand of the MTase domain central β-sheet and the loops between first and second and third and fourth strands

(Figure 3A). Structures of the NSP14/NSP10 complex from SARS-CoV in complex with S-adenosyl methionine (SAM), and with S-adenosyl homocysteine (SAH) and cap analogue GpppA^5^ show both substrates binding close together within the same large active site with the SAM/SAH binding and making interactions with the extended loop between 2^nd^ and 3^rd^ strands, whilst the methionine/homocysteine moiety approaches the GpppA in a manner that appears compatible with transfer of the methyl group to the Guanine N7.

**Figure 3.**
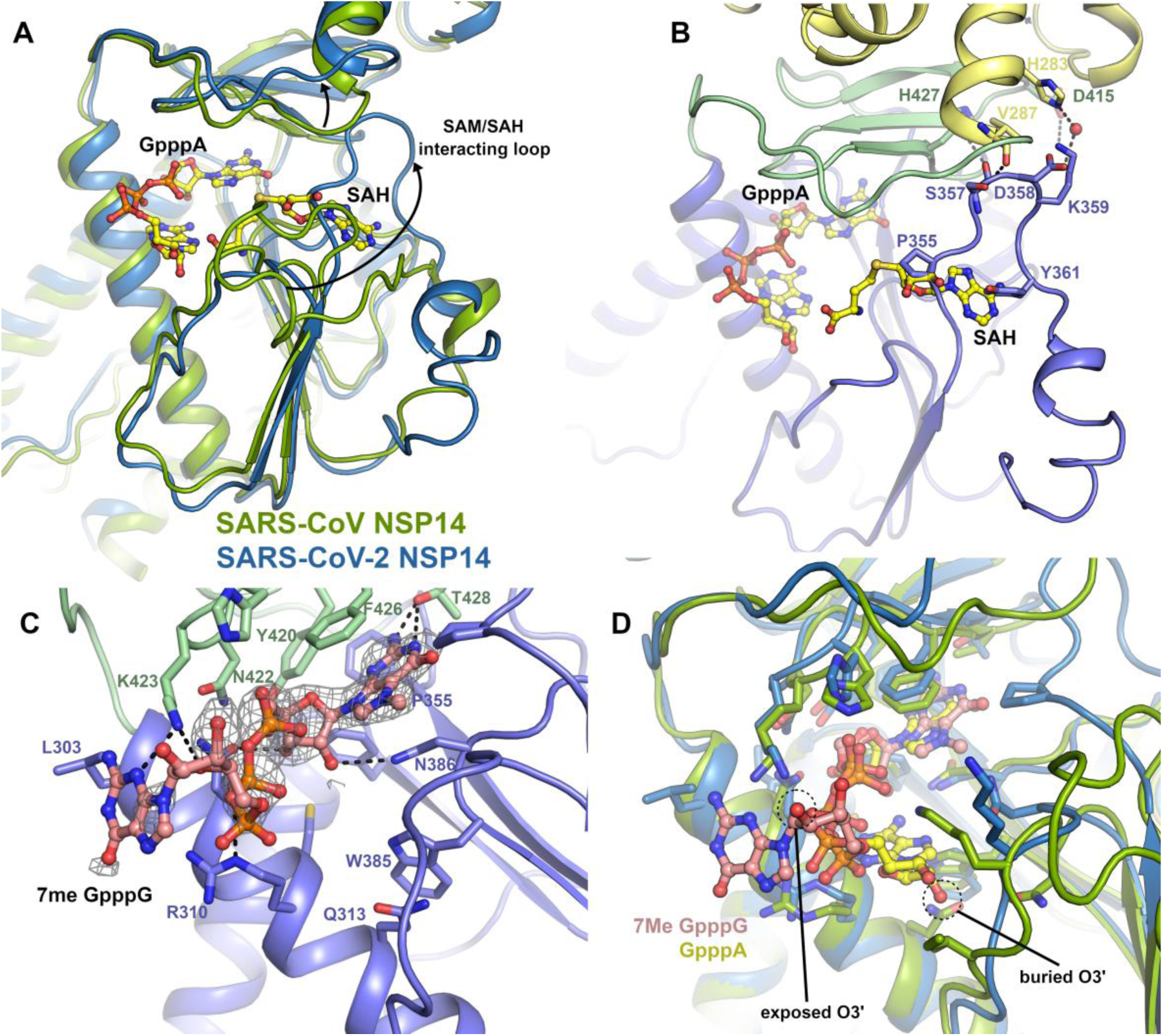
Structural differences in the NSP14 MTase active site. (**A**) Alignment of the SARS-CoV NSP14 GpppA/SAH ternary complex (shown in green) with the SARS-CoV-2 NSP14 MTase domain (shown in red), a significant conformational change can be seen for an extended loop that makes interactions with SAH in the SARS-CoV NSP14 structure. Both ligands are shown in stick format for reference. (**B**) Close up view of the SAM/SAH interacting loop in SARS-CoV-2 NSP14, residues which overlap with SAH or form contacts to residues in the ExoN or hinge domains are shown in the stick format. (**C**) Structure of SARS-CoV-2 NSP14 in complex with ^7Me^GpppG. The 2F°-1F^c^ electron density map is shown in grey in the vicinity of the ^7Me^GpppG contoured at 1σ. (**D**) Comparison of the binding poises of the GppA from SARS-CoV NSP14 with the ^7Me^GpppG from SARS-CoV-2 NSP14. The proteins are colored as for panel A and the positions of the O2’ which is the attachment point for full RNA cap substrates is marked.

Comparing these structures to our NSP14 MTase domain structure reveals generally good agreement (overall RMSD 0.8 Å) although large differences (displacements of up to 25 Å) can be seen in the conformation of the extended SAM/SAH interacting loop (residues 356-378). In our NSP14 structure, this loop makes several polar interactions with residues in the ExoN domain (S357 to V287, D358 water mediated to H283) and hinge domain (S577 to H427 and K359 to D415). The loop continues on the opposite side of the central β-sheet, in comparison to the SARS-CoV structures (Figure 3A). As was the case for regions of the N-terminal NSP10 interacting regions, several regions are observed interchange secondary structure elements with residues 363-366 forming a short section of helix rather than forming an additional sixth strand on the central MTase β-sheet. This loop conformation would not appear to be compatible with binding of SAM or SAH in the same manner as previously reported^5^ as the new loop conformation significantly overlaps with the adenine moiety. Consistent with this, attempts to soak either SAH or SAM to our crystals did not give any convincing electron density. On the other hand, the cap binding site appears to be open and we were able to soak in a cap analogue ^7Me^GpppG which gave clear electron density despite a slightly lower overall resolution (Figure 3B & S3). The ^7Me^GpppG has both methylated and un-methylated guanine moieties and as such is both a substrate and product mimic. The first nucleobase of SARS-CoV-2 transcripts is adenine although NSP14 has been demonstrated to be active on both GpppG and GpppA methyl cap analogues as substrates^23^.

The ^7Me^GpppG binds with the ^7Me^G in the product state with the ^7Me^G stacking between F426 and N386. The electron density map shows a clear bulge in the position of the methylated N7 (Figure S3) and the methyl group is buried fairly deep within the pocket and makes Van der Waals interactions with P355. The rest of the guanine moiety stacks against F426 and N386 and makes polar interactions with T428 and D388 in a similar manner to that described previously for SARS-CoV NSP14^5^. Other elements of the ^7Me^GpppG binding poise are less similar to previous structures, the second and third phosphates taking a slightly different path and forming polar contacts to K423 and R310. The ribose and nucleobase of the base 1 (adenine in the case of the SARS NSP14 but guanine for our structure) is significantly less well ordered in the electron density map (Figure S3) and is modeled in a position where it makes polar contacts to K423 (Figure 3C). The ribose O3 of the first nucleotide is in a solvent exposed position such that it would be possible to be incorporated into the NSP14 active site in the context of a full RNA cap molecule. This is in contrast to the SARS-CoV NSP14-GpppA-SAH ternary complex structure where the ribose is predominantly buried forming close contacts to K336 and I338. The discrepancy in the binding of the CAP analogues may be explained by the influence of crystal contacts in the SARS-CoV structure which would prevent the binding mode we have observed due to steric clashes (Figure S4). The SAH moiety in this structure also makes several steric clashes in the vicinity of the ribose and homocysteine moieties including clashes to the GpppA cap analogue itself (Figure 3D). Taken together, these observations suggest the potential for a more complex catalytic mechanism involving conformational changes to the flexible SAM/SAH binding loop and the possibility of an ordered sequential mechanism of substrate binding and product release.

### NSP14 as a constituent of the Replication Transcription Complex

A recent structure has been determined by cryo-EM for NSP14/NSP10 in complex with the rest of the RTC that has been dubbed Cap(0)-RTC^14^. This complex is comprised of NSP12, NSP7, NSP9, two copies of NSP8 and NSP13 and a single copy of the NSP14/NSP10 sub-complex. In this model NSP14/NSP10 primarily contacts NSP9 and the NiRAN domain of NSP12 via residues in the NSP14 ExoN domain. A subset (15%) of the particles picked from micrographs belong to a dimeric form which has additional contacts formed between the NSP14 MTase domain and the Zn-binding domain of NSP13 and the “fingers” and “thumb” region of the NSP12 polymerase^14^. In both monomeric and dimeric complexes, the conformation of the mobile SAM binding loop is identical to that of the previously reported SARS-CoV NSP14/NSP10 complex strucure^5^, although we note that the potential movements observed for this loop in our structure can be accommodated in both versions of the complex (Figure 4A, 4B). Outside of the MTase domain active site, the N-terminal NSP10 interacting region of the ExoN domain forms a significant overlap with residues in NSP9 (suggesting the interface is not compatible with NSP14 in the absence of NSP10) (Figure 4C). However, the recently published cryo-EM structure of NSP14/NSP10 in complex with RNA^9^ would lead us to question the possible biological significance of these complexes, due to the fact that the path of the RNA to the ExoN active site is completely blocked by the presence of NSP9 (Figure 4D) suggesting this particular complex is not compatible with exonuclease activity.

**Figure 4.**
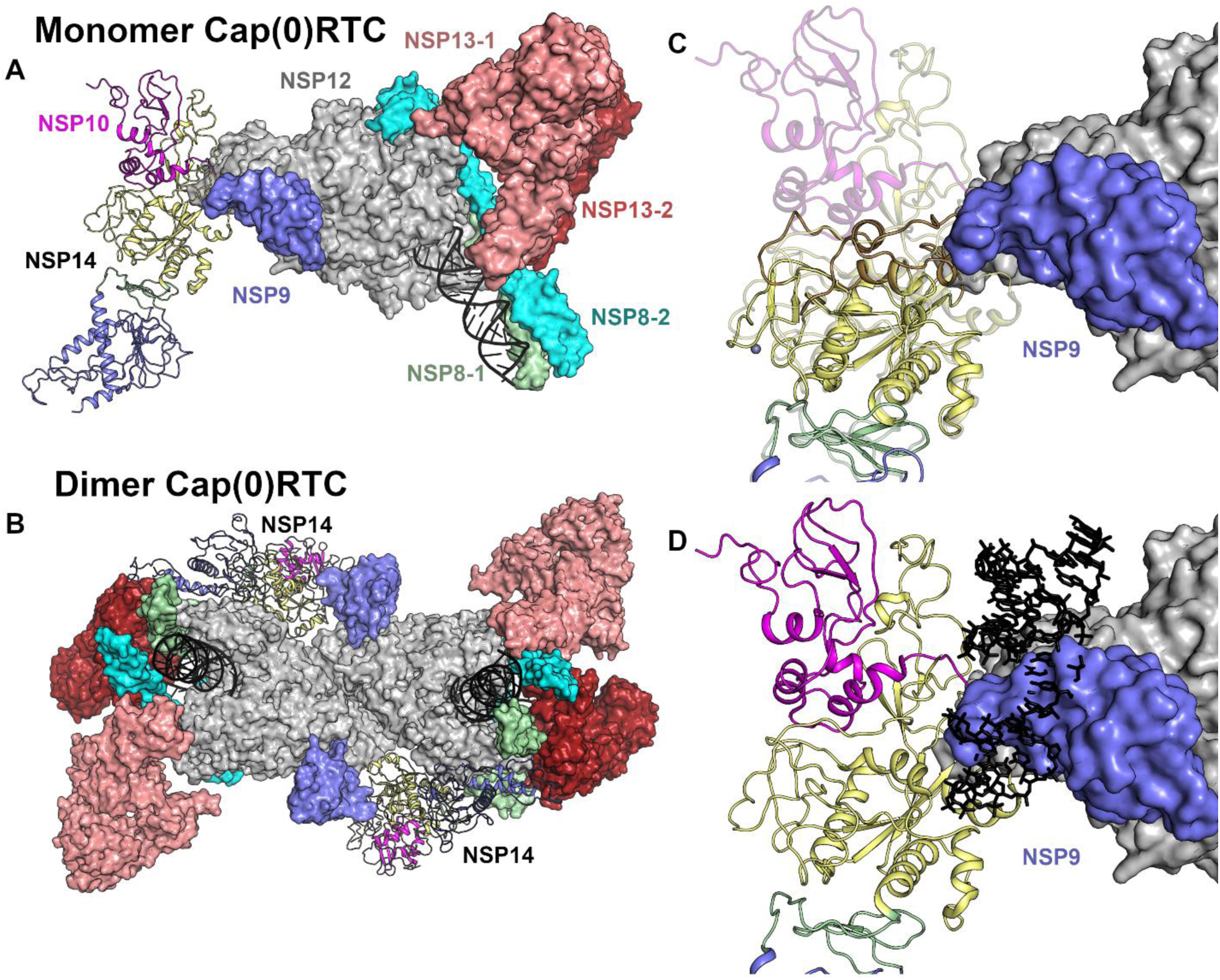
Evaluation of NSP14 as part of the Cap(0)RTC complex. (**A**) Overview of the positioning of NSP14 within the monomeric Cap(0)RTC complex based on the recently published Cryo EM structure^14^. Subunits are colored individually as labeled with NSP14 and NSP10 shown in the cartoon format. (**B**) Overview of the positioning of NSP14 within the Dimeric Cap(0)RTC complex as viewed along the two-fold symmetry axis. (**C**) Overlay of NSP14 alone with the NSP14 NSP10 complex within the Cap(0)RTC complex. The N-terminal region of NSP14 forms significant steric clashes with NSP9. (**D**) Overlay of the NSP14-NSP10-RNA complex (7N0B)^9^ onto the Cap(0)RTC complex. Severe steric clashes are evident with the RNA substrate (shown in black in stick format) suggesting this arrangement is not compatible with nuclease activity.

Moreover the Cap(0)-RTC complexes have been obtained using a modified form of NSP10 in which the proteolytic processing of the link to NSP9 had not occurred, based on the observation that other coronaviruses lacking a cleavage between NSP9 and NSP10 are viable^24^. The position of the NSP9 does not change when comparing complexes of the RTC with NSP9 alone^25^ or with the NSP9-NSP10/NSP14 fusion. This would suggest that the positioning of the fusion protein is dominated by the contacts made by NSP9. The authors of the Cap(0)-RTC do refer to a reconstruction without a NSP9-10 fusion that had “convincing density” for the NSP14, although this data has not been made available. The lack of other experimental evidence to support the model would lead us to suggest some caution in its interpretation.

### X-ray fragment screening of NSP14

The fact that our crystals capture NSP14 in a pre-activated state prior to its interaction with NSP10 represents a unique opportunity to exploit this for drug discovery. Our crystals grow relatively robustly, are tolerant to DMSO and standard soaking protocols and routinely diffract to around 2.0 Å resolution, sufficient for the reliable identification of binders. A total of 634 fragments were soaked using the DSI poised fragment library^26^ at a nominal ∼50 mM final concentration, yielding 588 diffraction datasets with the majority diffracting better than 2.5 Å. Analysis of the electron density maps using the PANDDA algorithm^27^ revealed a total of 72 fragments bound across 59 datasets (Figure 5A and supplementary Table I). The most prominent fragment hotspot was found in the MTase active site with 16 fragments (Figure 5B) with the majority of the fragments bound in the vicinity of the methylguanine and SAM moieties making stacking interactions with F426 and N386 and a number of polar contacts to nearby residues including N306, N334, N386, N388 and T428. No fragments were located in the vicinity of the ^7Me^GpppG phosphates although a smaller cluster of 3 fragments is bound close to the second guanine moiety and make polar contacts to Y296, N306 and N442 and hydrophobic interactions with P297 and I298. Another prominent pocket in the MTase domain with 11 fragments bound lies near the extended loop (residues 365-377) that was found to adopt a different conformation than in previous NSP14/10 complex structures. Fragments at this site make polar contacts to E365, L366 and N375 with a subset of the fragments inducing the disorder to order transition in loop 370-373 which is disordered in our crystals in the absence of fragment binding (Figure 5C). Whilst we note that a significant proportion of the pocket and interactions formed with these fragments are provided by residues in a crystal contact (Figure 5C), the site is a good candidate for allosteric inhibition due to the fragments interfering with the ability of the SAM interacting loop (residues 352-378) to transition to the conformation seen in the SAM bound NSP14/10 complex structures (Figure 5C).

**Figure 5.**
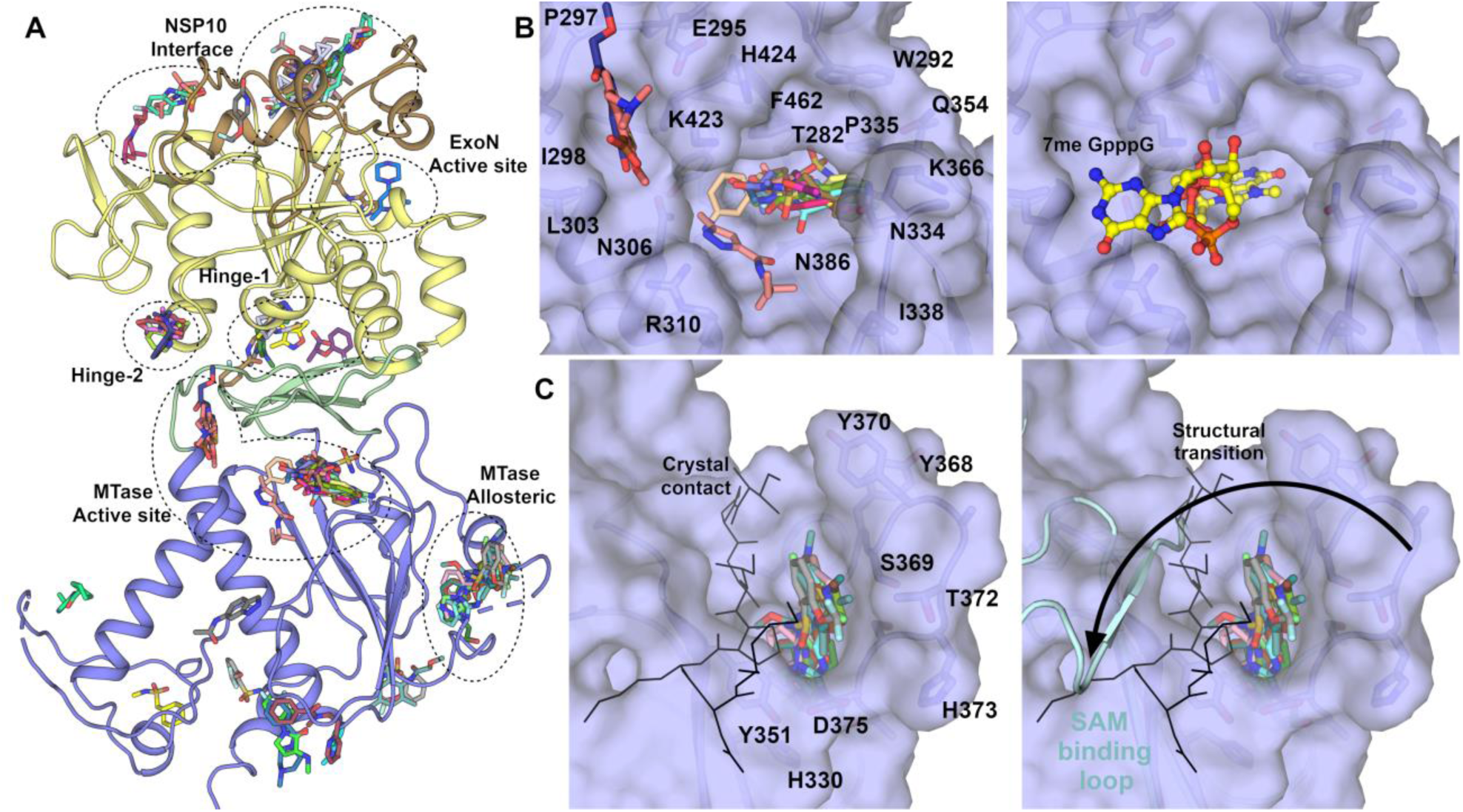
NSP14 fragment screening results overview and details of MTase domain pockets. (**A**) Overview of all 72 fragments observed to bind to NSP14 from our fragment screen. The major pockets and potential sites of inhibition are labelled. (**B**) Surface view of fragments bound to the MTase active site, the right hand panel shows the ^7Me^GpppG viewed from the same perspective for reference. (**C**) Surface view of pockets bound to a potential allosteric site of inhibition for the MTase activity. The black stick representation shows the contribution of a crystallographic symmetry mate to the upper left side of the pocket. The right hand panel shows the conformation of the equivalent loop (shown in Cyan) in the SARS-CoV NSP14/10 complex. This loop is involved in binding SAM, and the fragments may inhibit the MTase activity by preventing this transition.

Further potential exists for allosteric inhibition of Exonuclease activity by means of blocking the ability of NSP14 to interact with NSP10 (Figure 6A). Four pockets were found in regions which are on the NSP10 interaction interface or were observed to transition upon complex formation. The most extensive cluster contains 10 fragments which span a relatively broad pocket bounded on one side by the first and second helices at the N-terminus and the disordered loop 95-102 on the other. Fragments make predominantly hydrophobic interactions on one side with I42, P46 and M57 and on the other side more polar contacts to T103, L105, A119, W159 and K196 (Figure 6B). Three fragments in two pockets are bound directly upon the NSP10 interface (Figure 6C), and two fragments bind directly in the ExoN domain active site forming polar contacts to residues N108 and D273, the latter being involved in coordination of the catalytic metal ions (Figure 6D).

**Figure 6.**
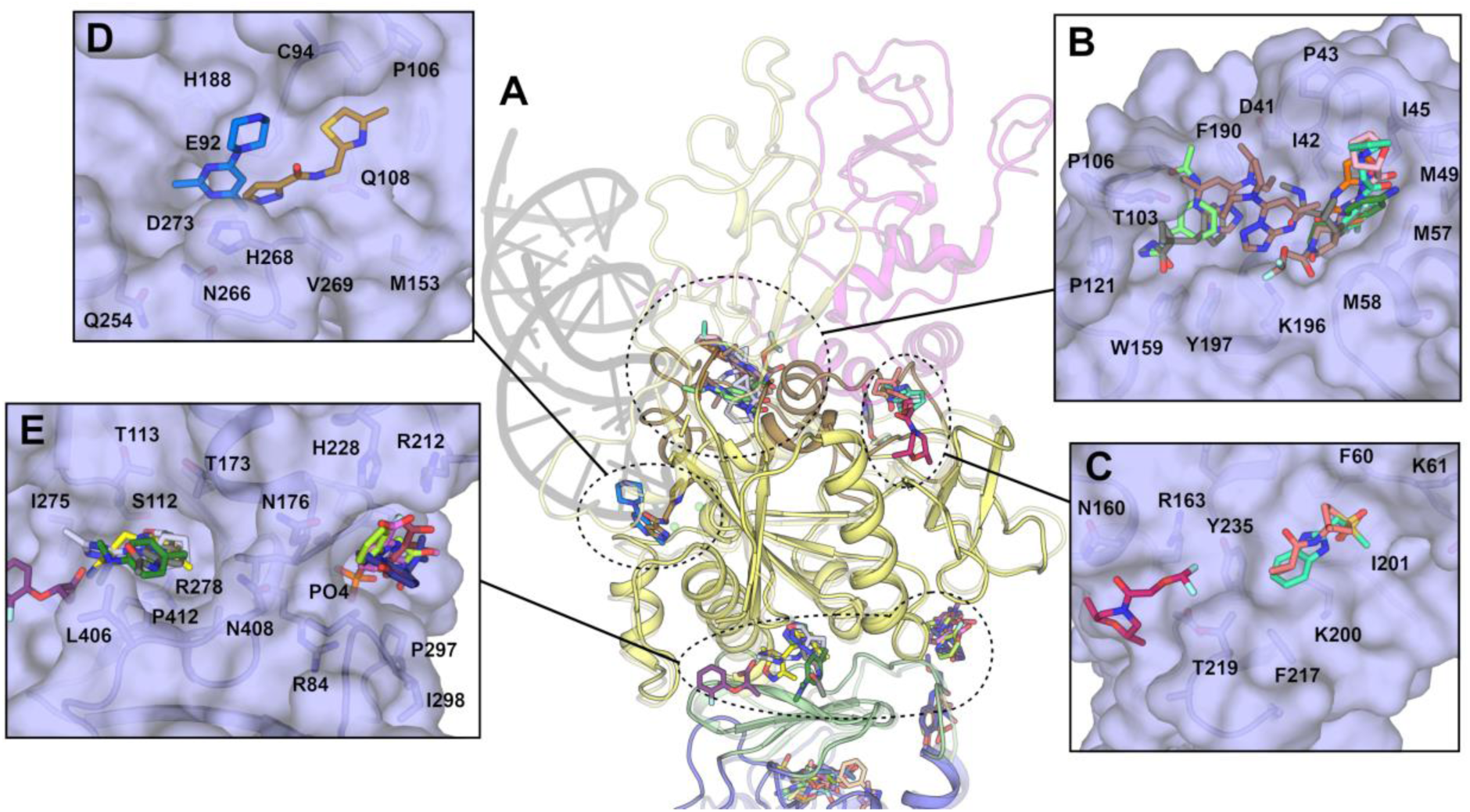
NSP14 fragments bound to pockets in the ExoN and hinge domains. (**A**) Overview of fragment binding sites with the NSP14/10 RNA complex shown in semitransparent representation for reference. (**B**) Surface view of fragments bound to NSP14 regions that undergo conformational changes to interact with NSP10. (**C**) Surface view of fragments bound directly on the NSP10 interface. (**D**) Surface view of fragments bound to the NSP14 ExoN active site. (**E**) Surface view of two clusters of fragments bound to pockets in the hinge domain.

Two clusters of fragments are observed to bind in the vicinity of the hinge domain that separates ExoN and MTase domains. The largest containing 6 fragments in a relatively deep pocket close to the ExoN domain Zn binding site. Fragments make polar contacts to S112 and H264, hydrophobic contacts to I275, L406 and P412 and numerous water mediated contacts (Figure 6E). The second cluster with 4 fragments bound lies directly opposite near a phosphate binding site in the ExoN domain. These fragments make hydrophobic contacts to P297 and I299 and polar contacts to R212, H228 and the Phosphate ion itself (Figure 6E). It is possible that the phosphate ion which makes quite extensive interactions at this site may play a role in RNA substrate recognition, furthermore the residue P297 which lines this pocket was identified as being essential for ExoN activity in SARS-CoV NSP14^16^. The existence of such mutants suggests that it may be possible to develop inhibitors that inhibit both enzymatic activities.

## Summary

In this study we have determined the crystal structure of SARS-CoV-2 NSP14 in the absence of its partner NSP10 to high resolution. NSP14 is an important target for development of possible antivirals for several reasons. Direct inhibition of the nuclease proofreading activity may increase susceptibility of SARS-CoV-2 to nucleotide inhibitors such as remdesivir^28,29^. Our structure offers a unique opportunity for discovery of inhibitors that bind to the ExoN domain in its inactive state with the potential to block the interaction with NSP10. The MTase domain is also a promising site to target for broad spectrum coronavirus inhibition, and potentially druggable pockets have been identified that are well conserved across coronavirus species^30^. To this end, potent SAM analogue inhibitors have been developed against NSP14 that show mid nM potency and promising selectivity profiles^31,32^. Our fragment screens identify 72 fragments bound in several promiscuous pockets that represent useful starting points for inhibitor development or may be used for the structure based elaboration of existing hits. It is important to note that these fragments are soaked into crystals at relatively high concentrations and whilst they may have good ligand efficiency, are not expected to be potent inhibitors without further optimization. We suggest the druggable MTase domain active site with its large number of fragment hits, and high degree of sequence conservation may be the best place to start for the development of anti-viral therapeutics that may be able to combat the current pandemic and also future emerging viral threats.

## Methods

### Cloning and expression of NSP14

The plasmid for NSP14 with His_6_ and Z-Basic tags at the N-terminal was synthesized in a pNIC-ZB vector with codon optimization for expression in *Escherichia coli* (Supplementary Table 1). The plasmid was transformed into BL21(DE3)-RR-pRARE competent *E. coli*. Cell cultures were grown in Terrific Broth media (Merck) supplemented with 20 μM zinc chloride at 37 °C, shaking at 180 rpm. Once OD_600_ reached 2.5, IPTG was added to the media at a final concentration of 300 μM. Cultures were incubated overnight at 18 °C, shaking at 180 rpm. Cells were harvested by centrifugation at 4200 rpm at 4 °C for 25 min.

### Purification of NSP14 protein

Cells were re-suspended in lysis buffer (50 mM HEPES pH:7.5, 500 mM NaCl, 10 mM Imidazole, 5% Glycerol, 1 mM TCEP) supplemented with protease inhibitor (Merck Protease inhibitor cocktail III, 1:1000) and 1X benzonase nuclease. Cells were sonicated for 20 min (5 sec pulse-on, 10 sec pulse-off). Cell debris was removed by centrifugation with Beckman JA-17 rotor at 17000 rpm at 4 °C for 45 min. The supernatant was incubated with 7 ml pre-equilibrated Ni-IDA resin in tubes at 4 °C for 2 hours. After batch binding, the tubes were spun at 700 × g at 4 °C for 5 min and the supernatant was discarded. Resin was washed with 30 ml lysis buffer twice and transferred into a gravity column with 25 ml W20 buffer (50 mM HEPES pH:7.5, 500 mM NaCl, 20 mM Imidazole, 5% Glycerol, 1 mM TCEP). Resin was washed again with 25 ml W30 buffer (50 mM HEPES pH:7.5, 500 mM NaCl, 30 mM Imidazole, 5% Glycerol, 1 mM TCEP). Protein was eluted with 25 ml IMAC elution buffer (50 mM HEPES pH:7.5, 500 mM NaCl, 300 mM Imidazole, 5% Glycerol, 1 mM TCEP) twice. Elution fractions were pooled and applied onto a pre-equilibrated 5 ml-HiTrap-SP High Performance column using a syringe. Column was washed with 50 ml SP wash buffer (50 mM HEPES pH:7.5, 500 mM NaCl, 5% Glycerol, 1 mM TCEP). Protein was eluted with 25 ml SP elution buffer (50 mM HEPES pH:7.5, 1 M NaCl, 5% Glycerol, 1 mM TCEP). Elution fraction was diluted in 50 mM HEPES pH:7.5, 5% Glycerol, 1 mM TCEP to reduce salt concentration to 500 mM before TEV cleavage. Diluted sample was treated with TEV protease (1:5 mass ratio) for 3.5 hours at room temperature. To remove ZB tag, the cleavage sample was passed through a 5 ml-HiTrap-SP HP column and the flow-through was collected. The flow-through fraction was concentrated with 30 MWCO Amicon filters and loaded onto a HiLoad 16/600 Superdex 200 pg column pre-equilibrated with SEC buffer (50 mM HEPES pH:7.5, 500 mM NaCl, 5% Glycerol, 1 mM TCEP). Pure NSP14 fractions were collected concentrated and stored at -80 °C for future experiments.

### Crystallization

Crystallization was performed by sitting-drop vapor-diffusion method with SGC-HIN3 HT-96 screen (Molecular Dimensions). Crystal plates were set up with 10 mg/ml protein and HIN3 screen, using 150 nl drops (1:1 ratio). Rod shaped crystals were grown in condition containing 1.26 M sodium phosphate monobasic, 0.14 M potassium phosphate dibasic at 4 °C. Further crystals were grown in this single condition from Molecular Dimensions. For NSP14-^7Me^GpppG crystals, 5 mM of m^7^GP_3_G monomethylated cap analogue (Jena Bioscience, NU-852) was soaked into mature NSP14 crystals (grown in 1.26 M sodium phosphate monobasic, 0.14 M potassium phosphate dibasic) overnight at 4 °C. Crystals were cryo-protected in a solution consisting of well solution supplemented with 20 % Glycerol, loop-mounted, and flash-frozen in liquid nitrogen.

### Structure Determination

All data were collected at Diamond light source beamlines I03 and I04-1 processed using XDS^33^. Data were moderately anisotropic and were corrected using an anisotropic cut-off as implemented in the STARANISO server^34^, which gave a significant improvement in the quality and interpretability of electron density maps. The structures were solved by molecular replacement using the program PHASER and the structure of SARS-CoV-1 NSP14 (5C8S) as a search model. Refinement was performed using PHENIX REFINE^35^. A summary of the data collection and refinement statistics are shown in table 1.

### X-ray Fragment Screening

A total of 634 fragments from the DSI poised fragment library (500 mM stock concentration dissolved in DMSO) were transferred directly to NSP14 crystallization drops using an ECHO liquid handler (50 mM nominal final concentration), and soaked for 1–3 h before being loop mounted and flash cooled in liquid nitrogen. A total of 588 datasets were collected at Diamond light source beamline I04-1 and processed using the automated XChem Explorer pipeline^36^. Structures were solved by difference Fourier synthesis using the XChem Explorer pipeline. Fragment hits were identified using the PanDDA program^27^.

Refinement was performed using BUSTER. A summary of data collection and refinement statistics for all fragment bound datasets is shown in Supplementary Table 3 and an overview of electron density maps for each hit is shown in Supplementary Table 2.

## Acknowledgements

The crystallographic screen was supported by the XChem facility at Diamond Light Source. We thank all the staff of Diamond Light Source beam-lines I03, I04 and I04-1 for providing support under Proposals LB26998, LB22717 and MX28172. This work was supported by a NCI P01 CA092584 grant and the Cancer Research UK Programme Award A24759.

## Author Contributions

J.A.N. and Y.Y. initiated the project. Y.Y., J.A.N. and N.I. performed expression, protein purification, and crystallization. N.I. and J.A.N. performed crystal optimization and fragment soaking, Crystal mounting, and XChem data management. J.A.N. performed X-ray data analysis and review of fragment binding.

J.A.N. and N.I. wrote the original draft manuscript. All authors read and approved the manuscript.

## Competing interests

The authors declare no competing interests.

## Data availability

Crystallographic coordinates and structure factors for all structures have been deposited in the Protein Data Bank with the following accessing codes: 7QGI, 7QGF, 5KSW, 5SKX, 5SKY, 5SKZ, 5SL0, 5SL1, 5SL2, 5SL3, 5SL4, 5SL5, 5SL6, 5SL7, 5SL8, 5SL9, 5SLA, 5SLB, 5SLC, 5SLD, 5SLE, 5SLF, 5SLG, 5SLH, 5SLI, 5SLJ, 5SLK, 5SLL, 5SLM, 5SLN, 5SLO, 5SLP, 5SLQ, 5SLR, 5SLS, 5SLT, 5SLU, 5SLV, 5SLW, 5SLX, 5SLY, 5SLZ, 5SM0, 5SM1, 5SM2, 5SM3, 5SM4, 5SM5, 5SM6, 5SM7, 5SM8, 5SM9, 5SMA, 5SMB, 5SMC, 5SMD, 5SME, 5SMF, 5SMG, 5SMH, 5SMI.

## Supplementary Information for

**Figure S1.**
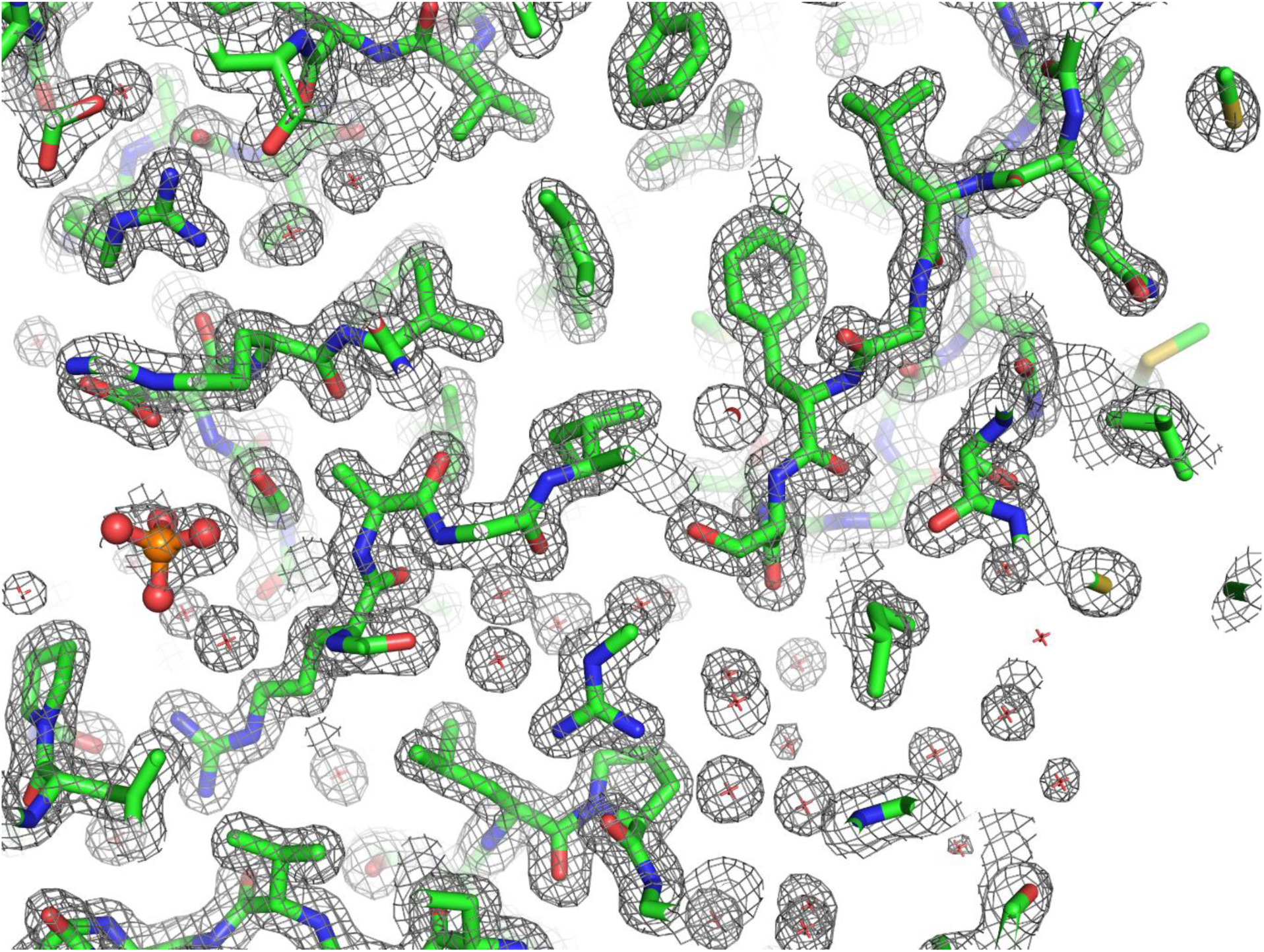
Representative 2F_o_-1F_c_ electron density map of NSP14 contoured at 1.5 σ.

**Figure S2.**
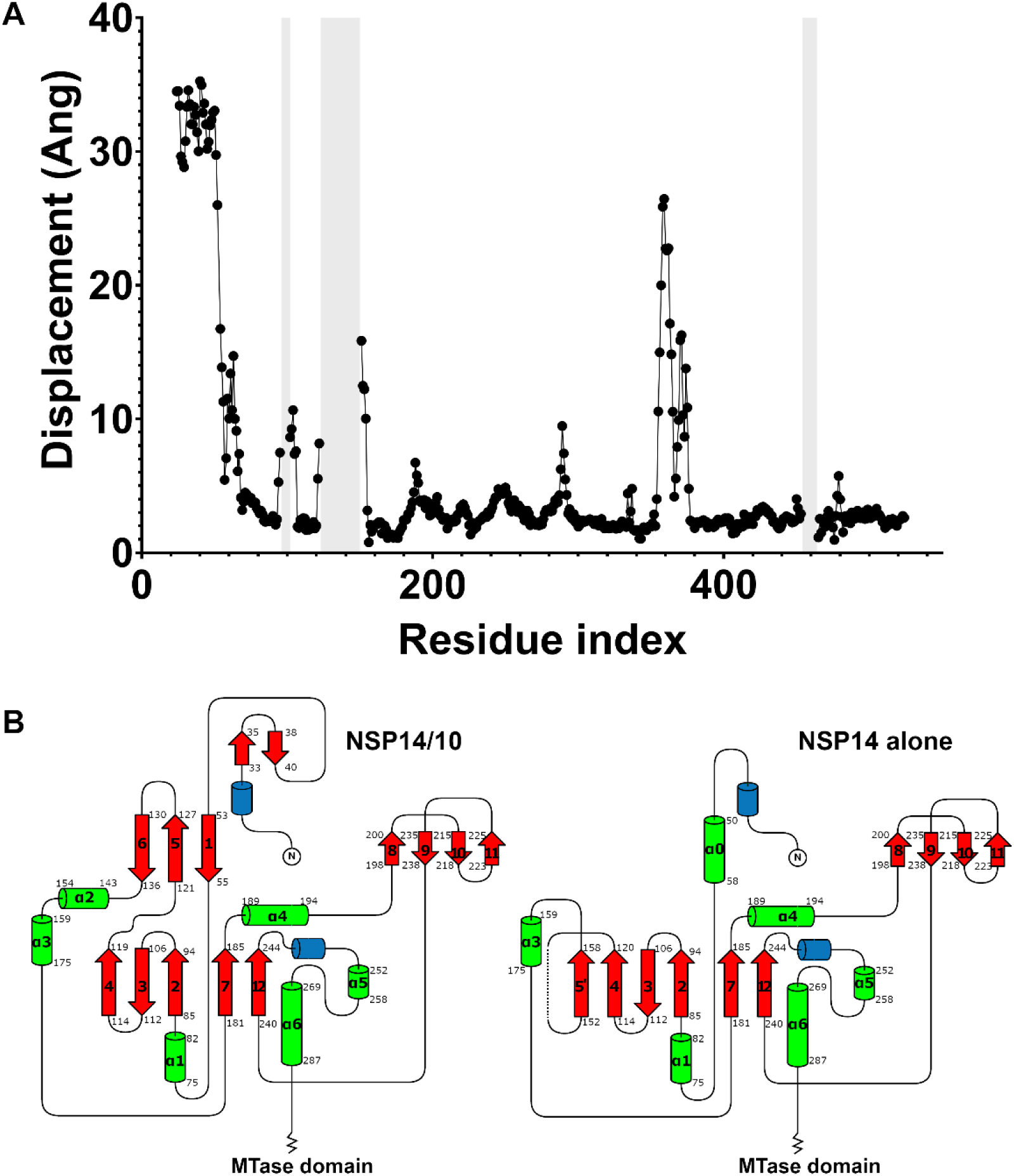
Comparison of NSP14 alone with NSP14 in the NSP14/NSP10 complex. (**A**) Cα displacements in Angstrom plotted as a function of residue number. The shaded grey areas represent regions disordered in the NSP14 structure. (**B**) Topology diagram showing the changes in secondary structure elements between NSP14/NSP10 complex and NSP14 alone. For consistency the nomenclature of the complex has been applied to the NSP14 alone structure.

**Figure S3.**
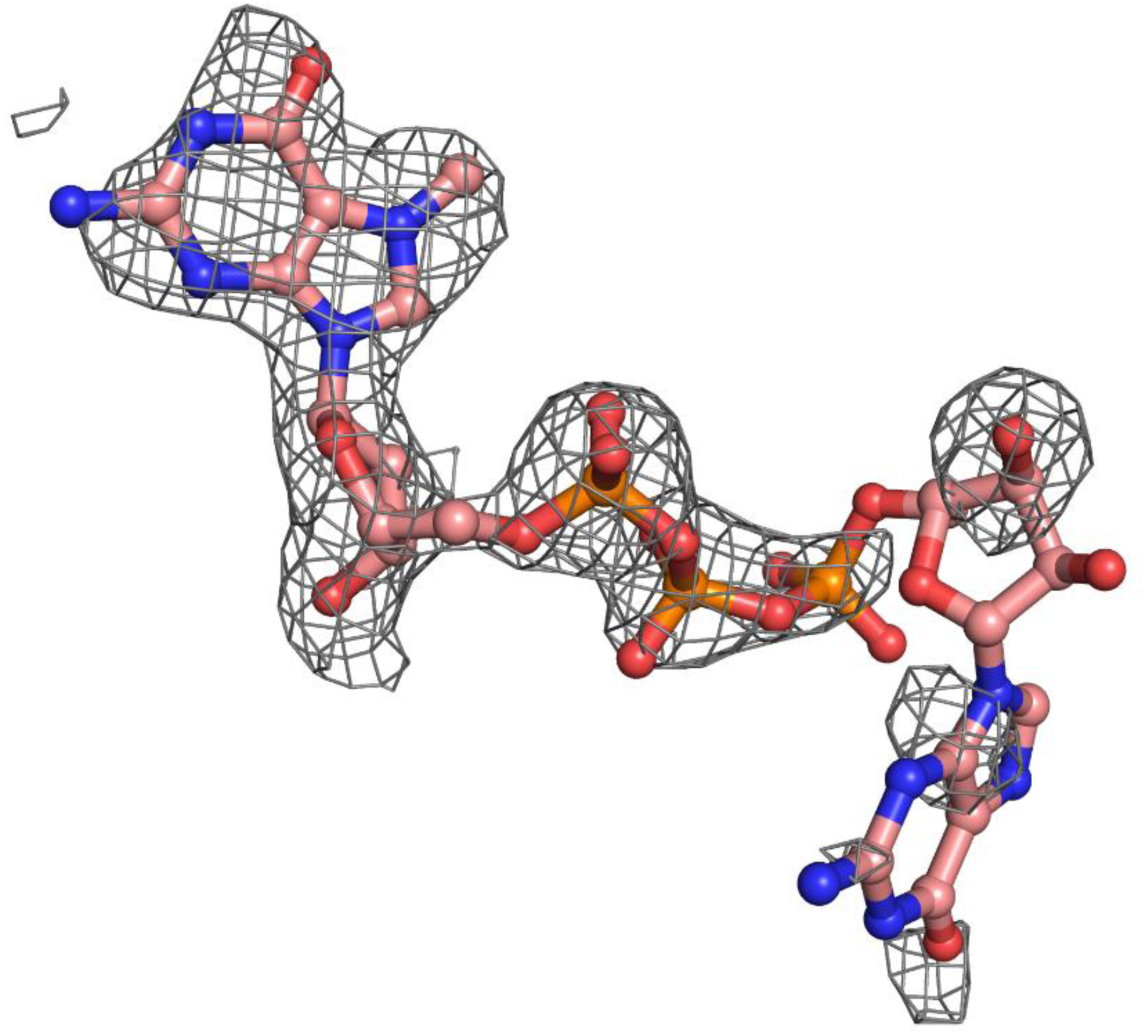
2F_o_-1F_c_ electron density map at 2.5 Å resolution contoured at 1σ in the vicinity of the ^7Me^GpppG. The methyl bulge is clearly visible in the electron density whilst the ribose and Guanine of the base 1 are less well ordered.

**Figure S4.**
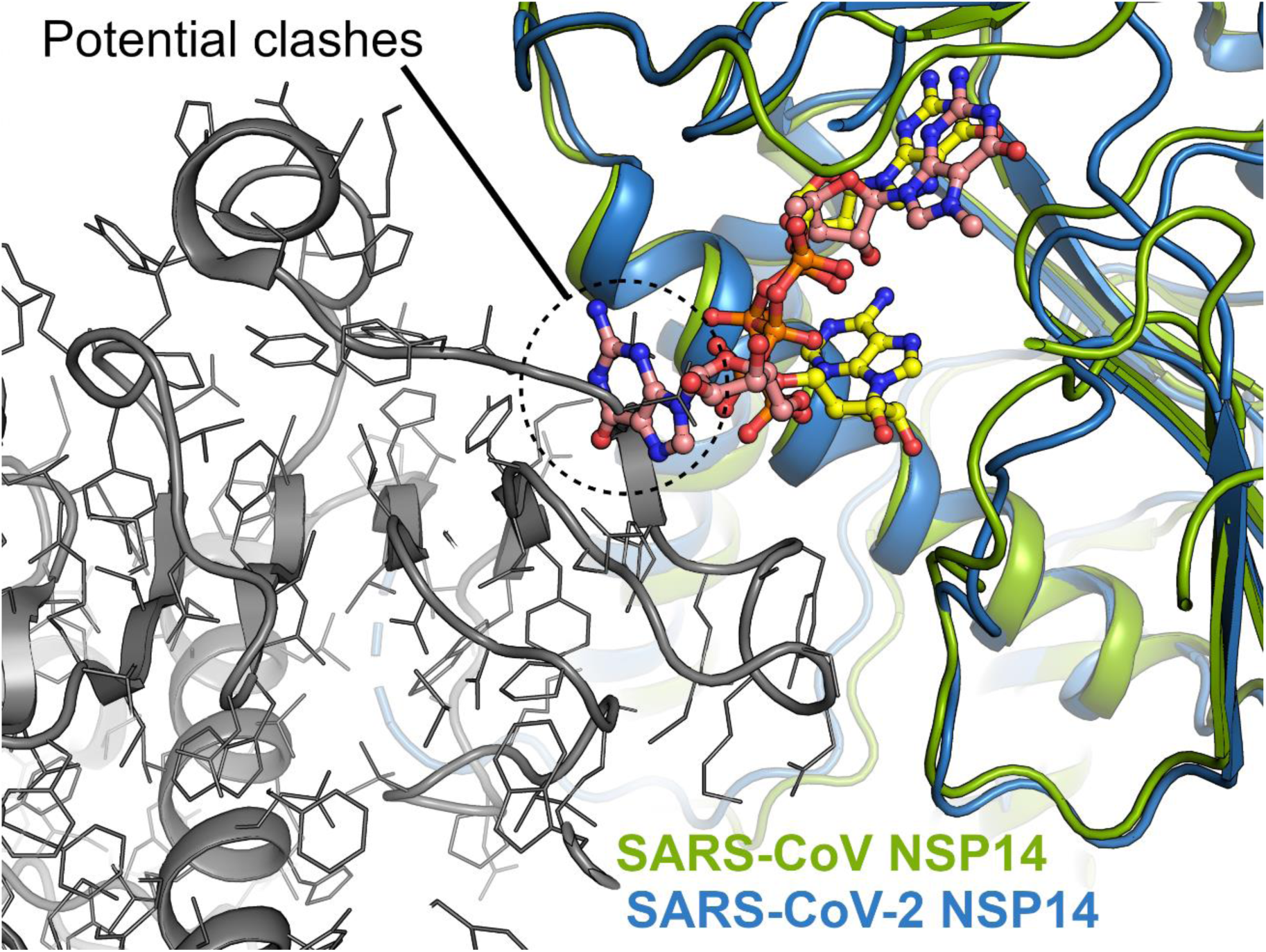
Comparison of SARS-CoV-2 NSP14 (Blue with pink ^7Me^GpppG) with SARS-CoV NSP14 (Green with yellow GpppA). The symmetry mate of SARS-CoV NSP14 (shown in grey) approaches close to the free ribose O3’ and would appear to prevent the ^7Me^GpppG conformation being adopted in the SARS-CoV-2 NSP14 due to steric clashes.

**Supplementary Table 1:**
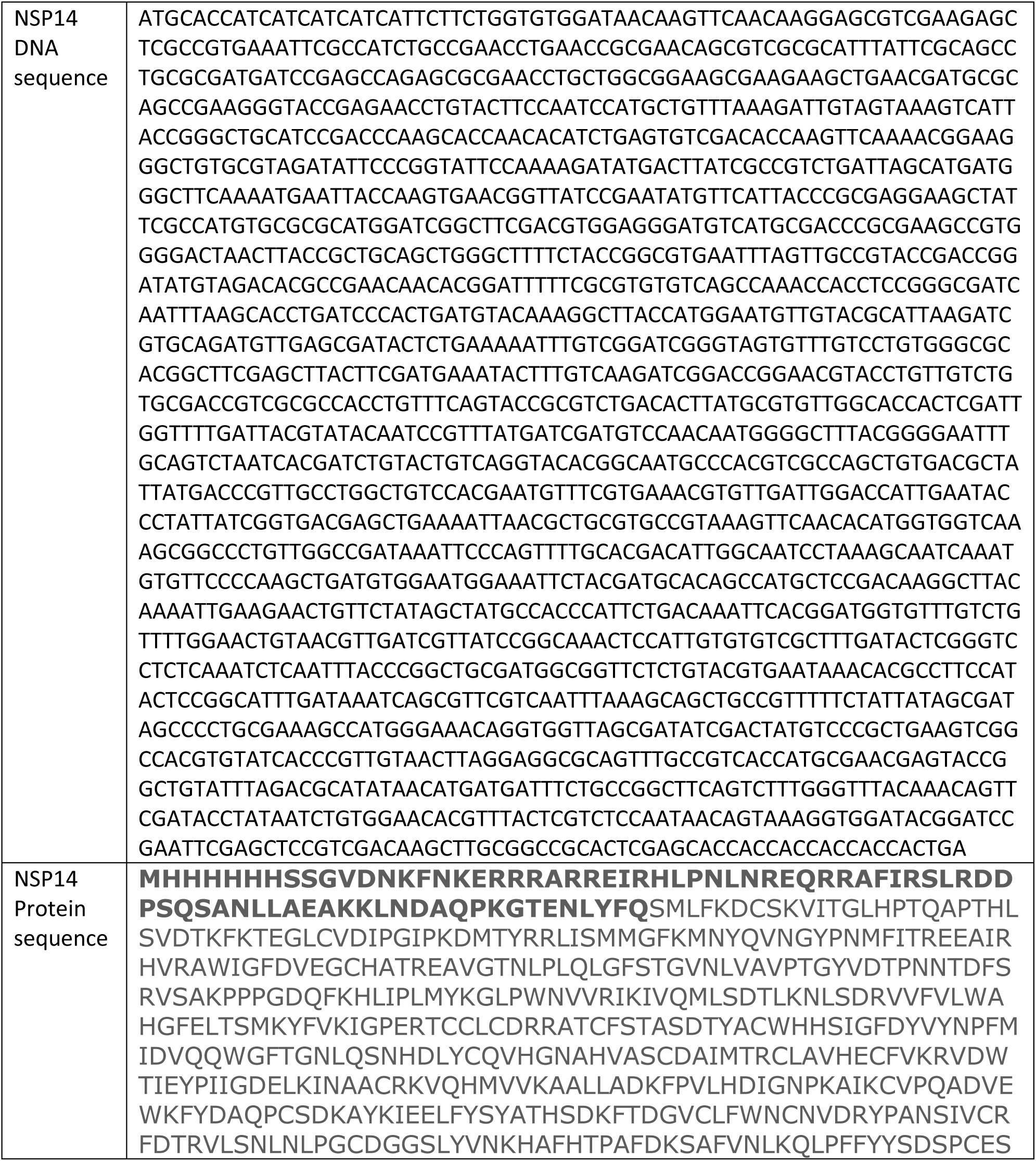
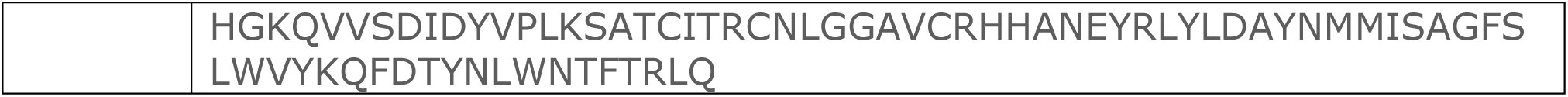
DNA and protein sequences for the codon optimized NSP14 construct used in this study. The bold sequence is removed during purification by TEV protease cleavage.

**Supplementary Table 2.**
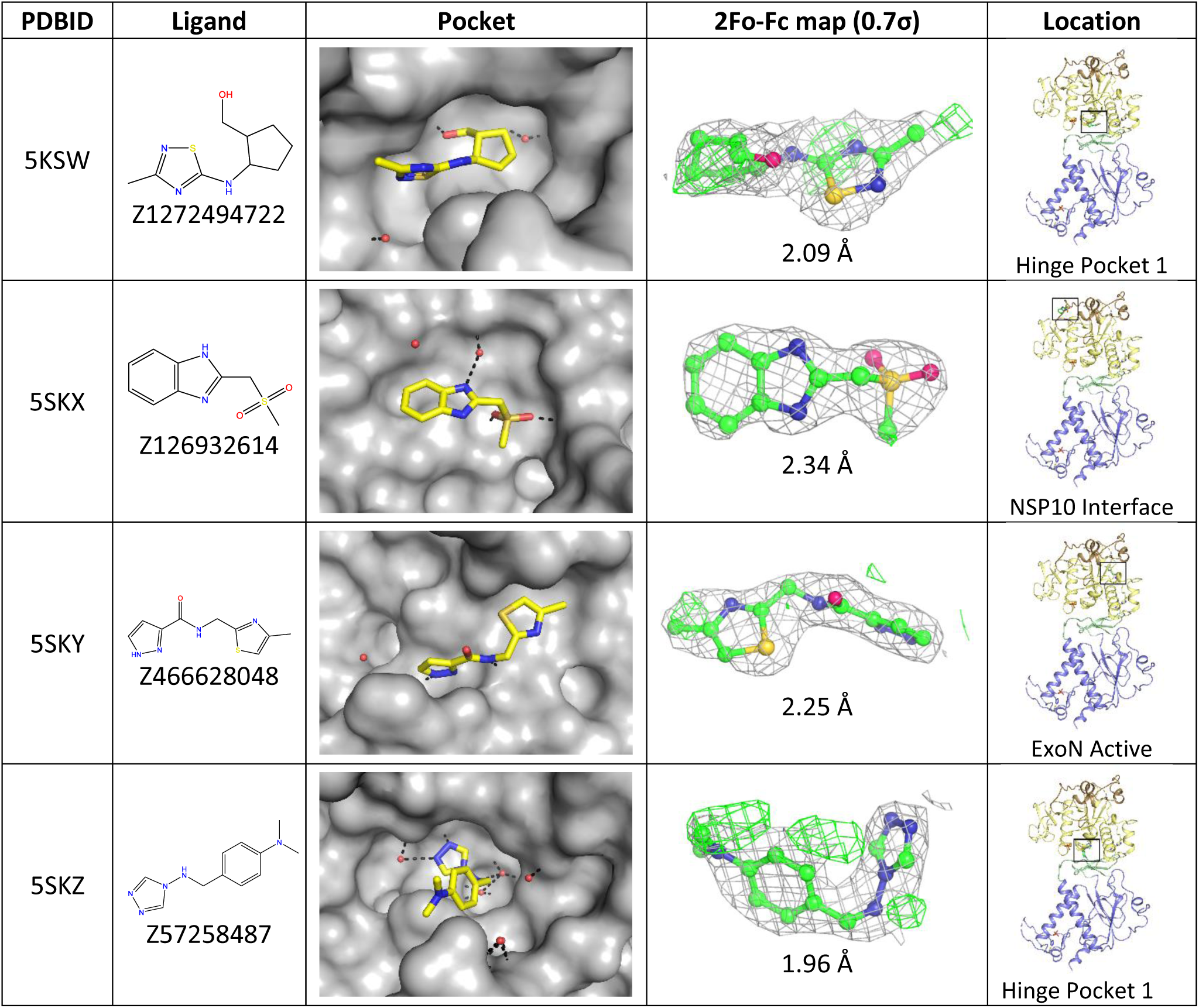
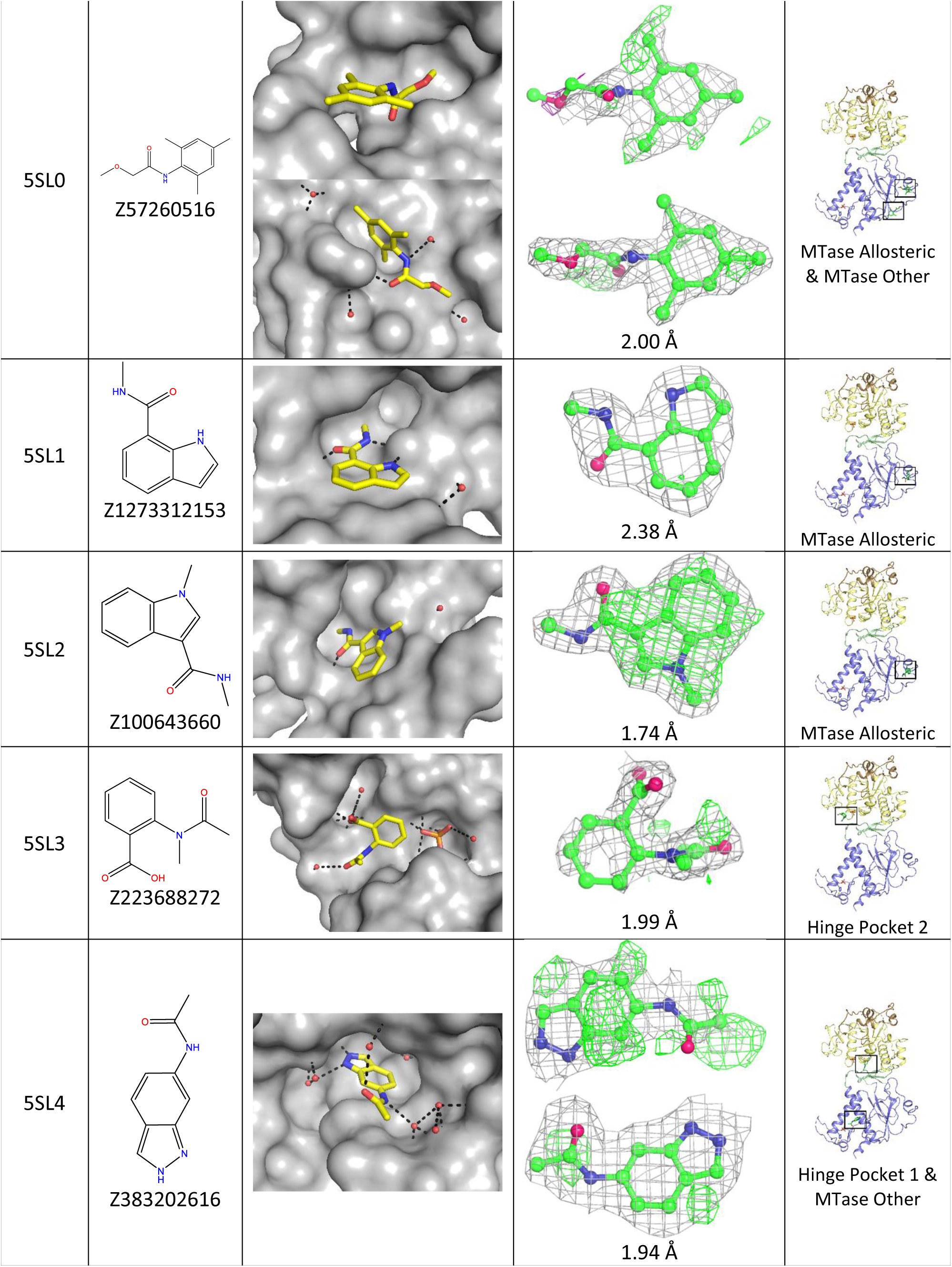
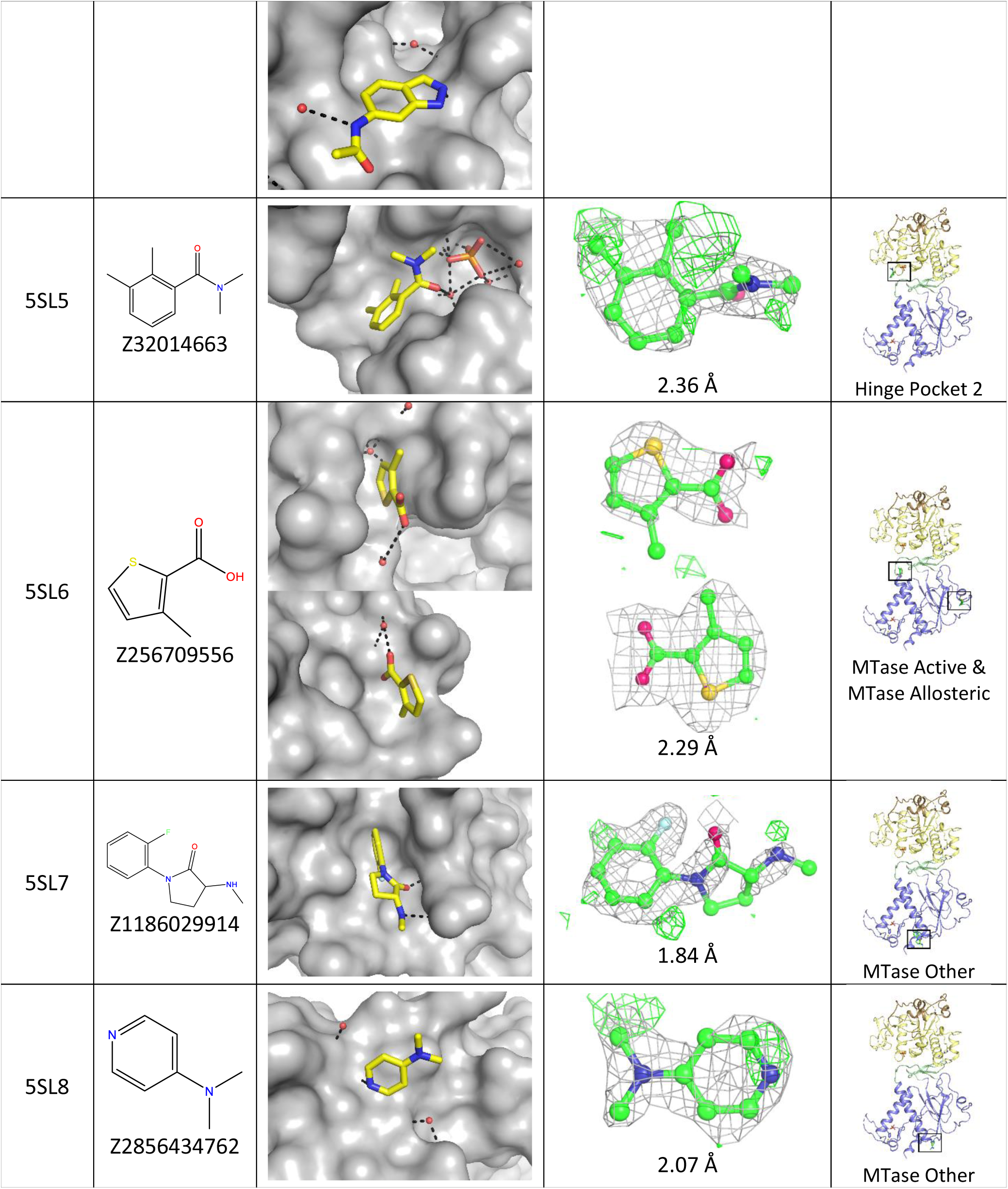
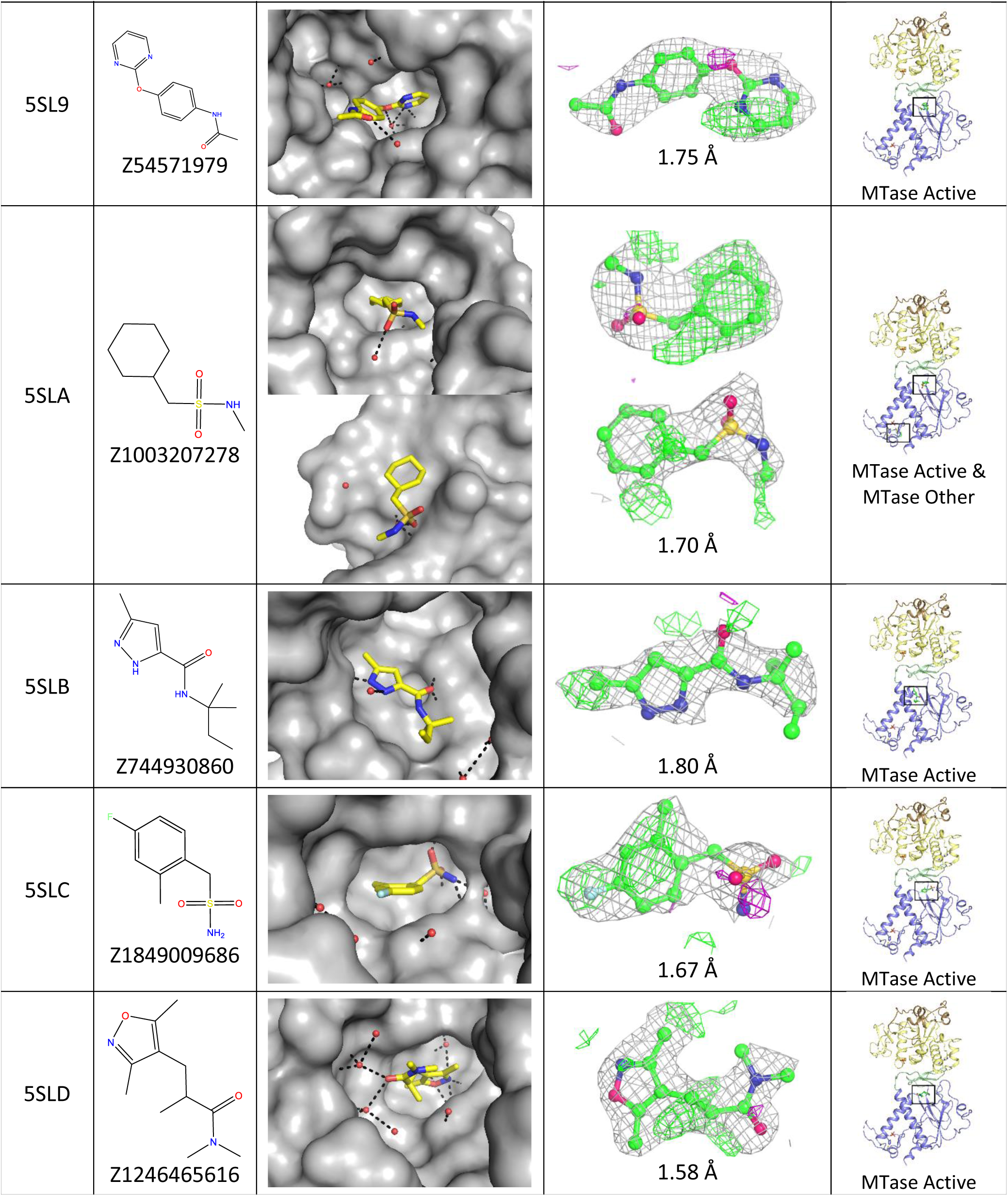
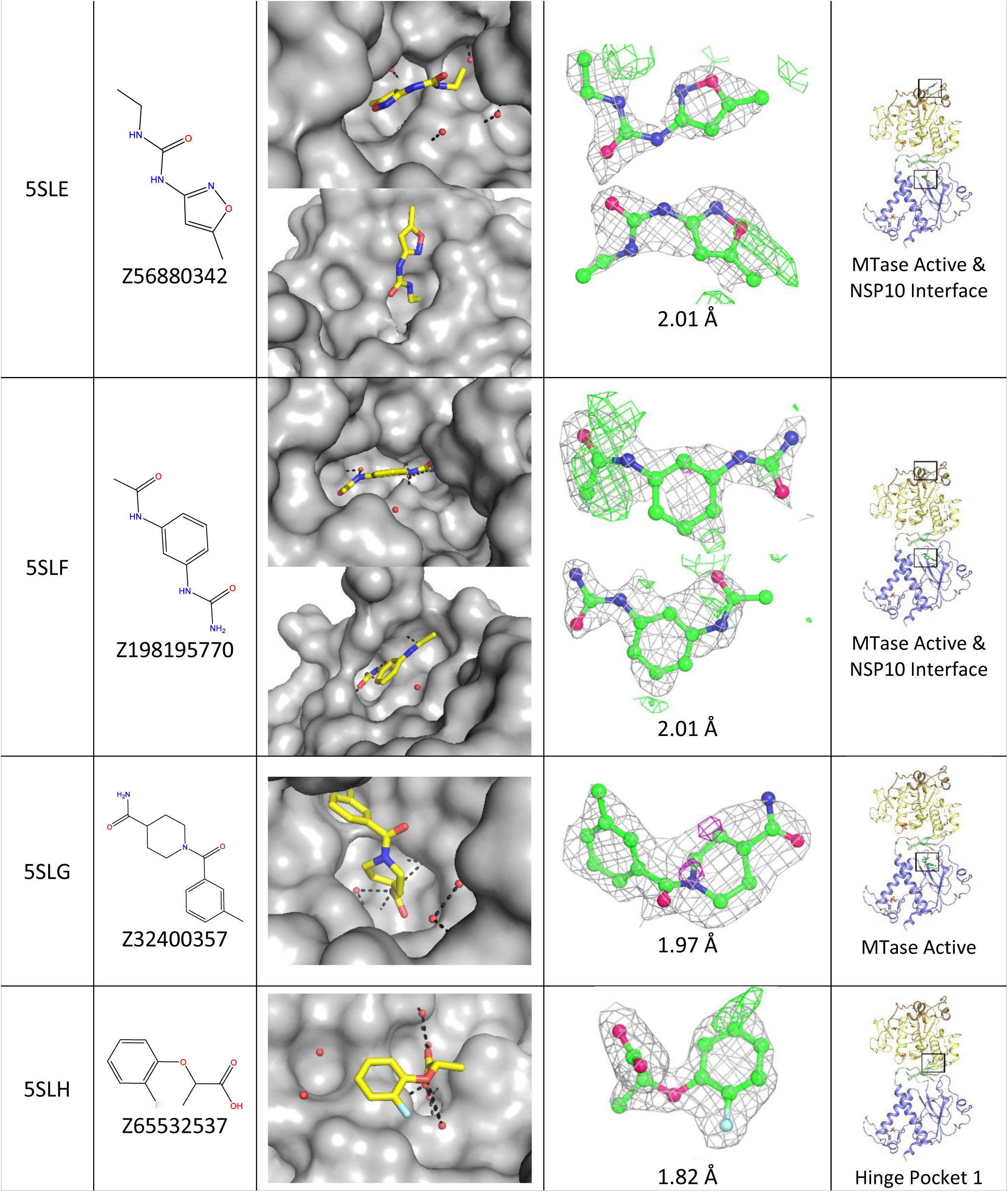
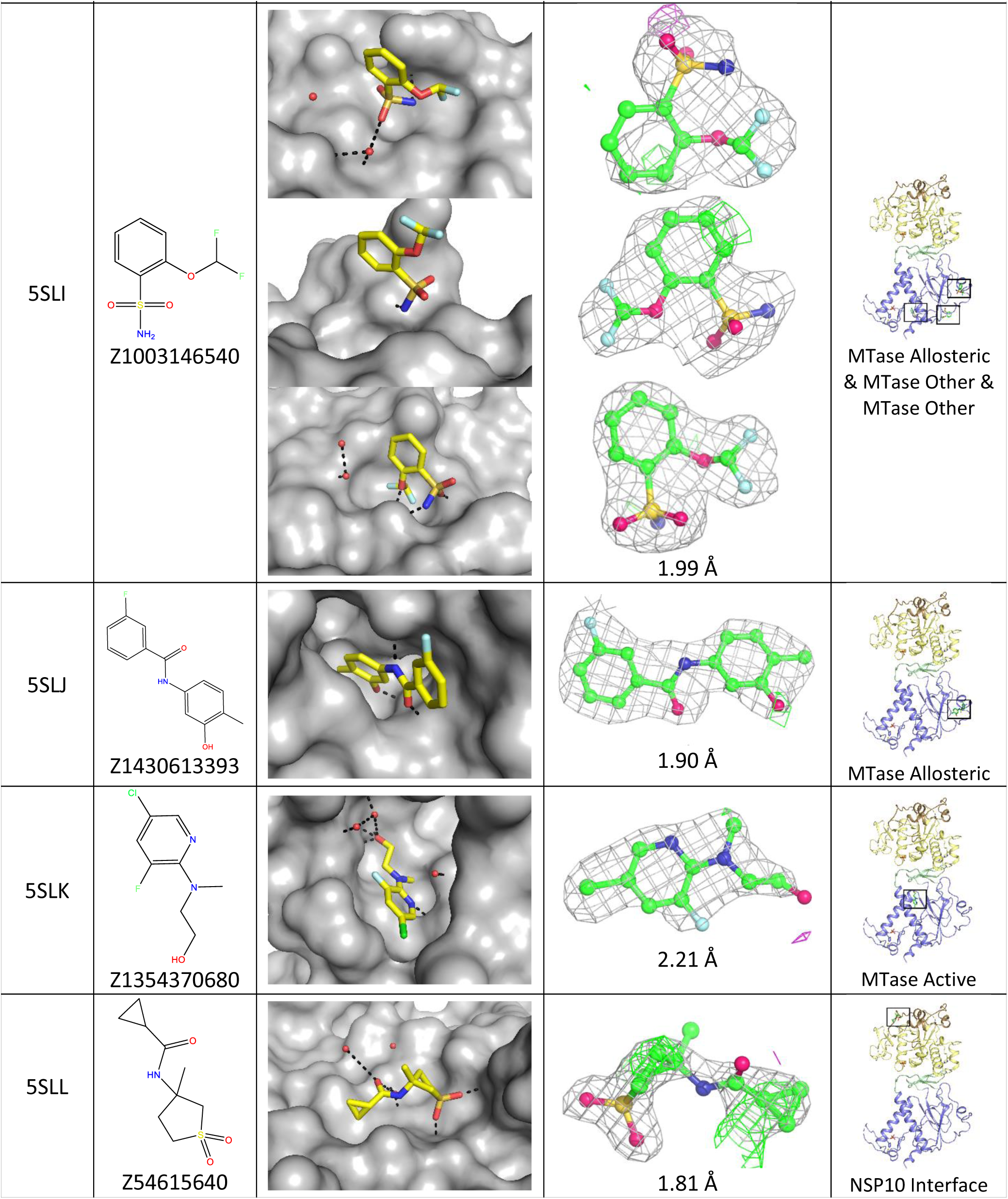
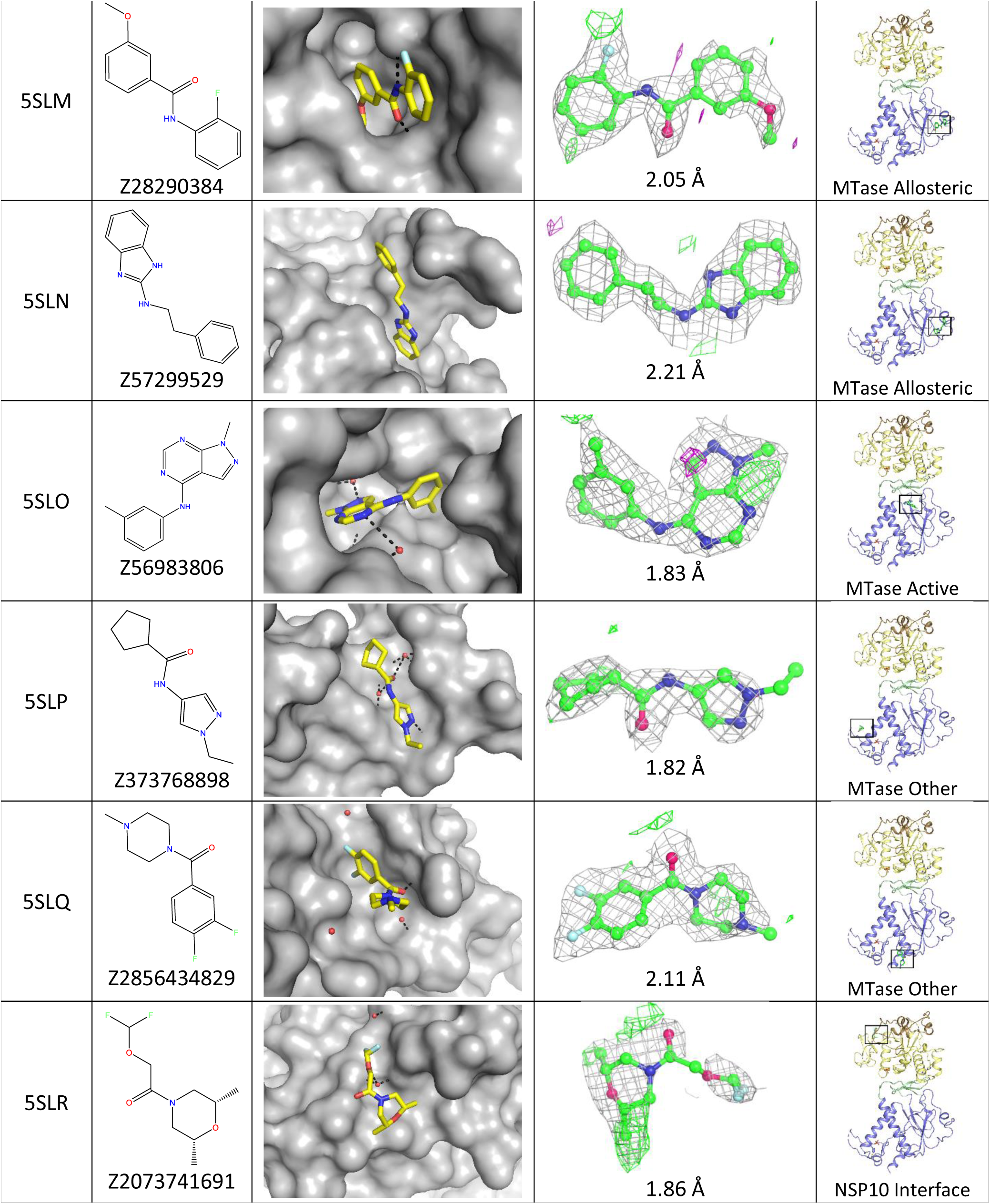
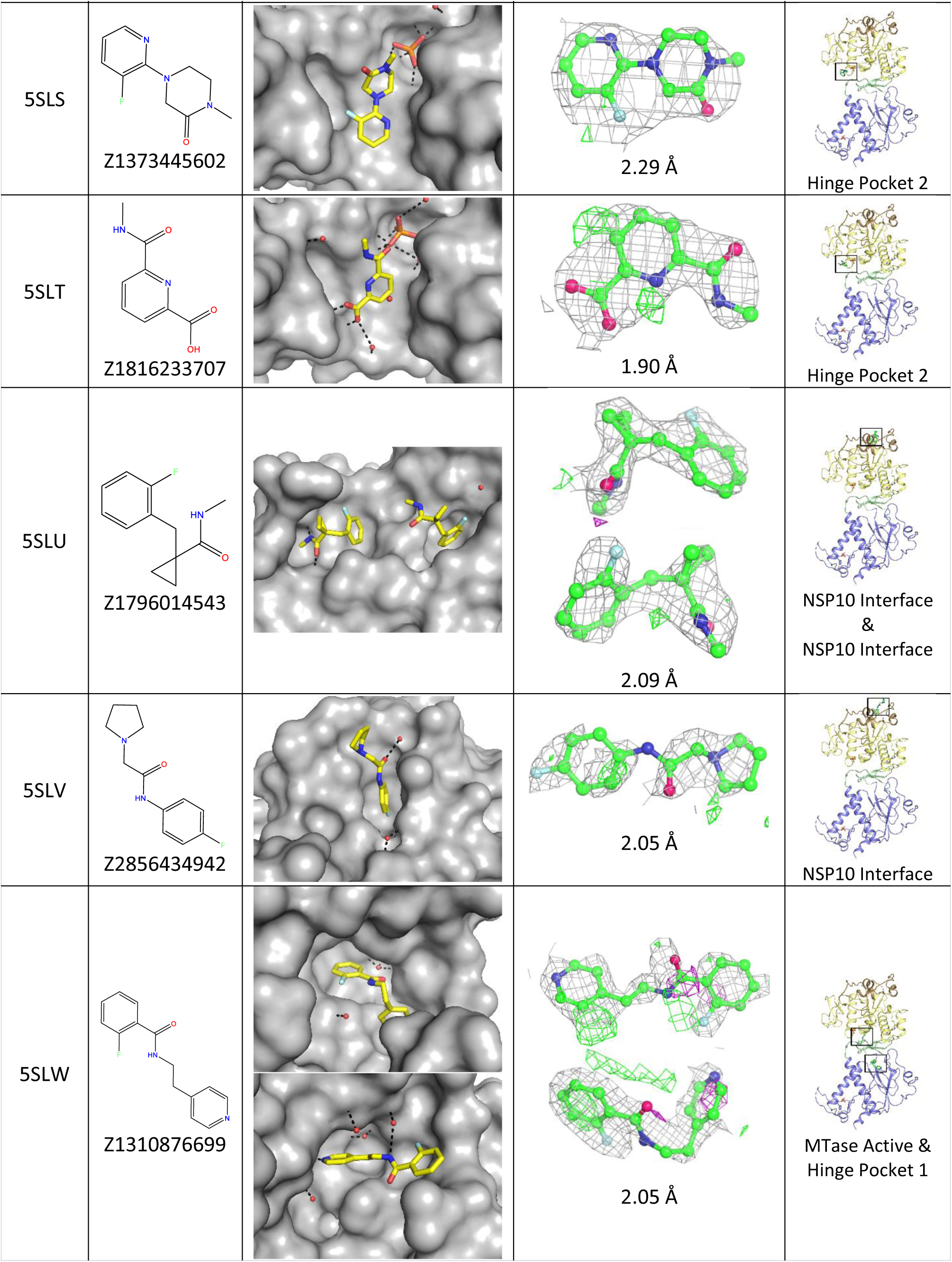
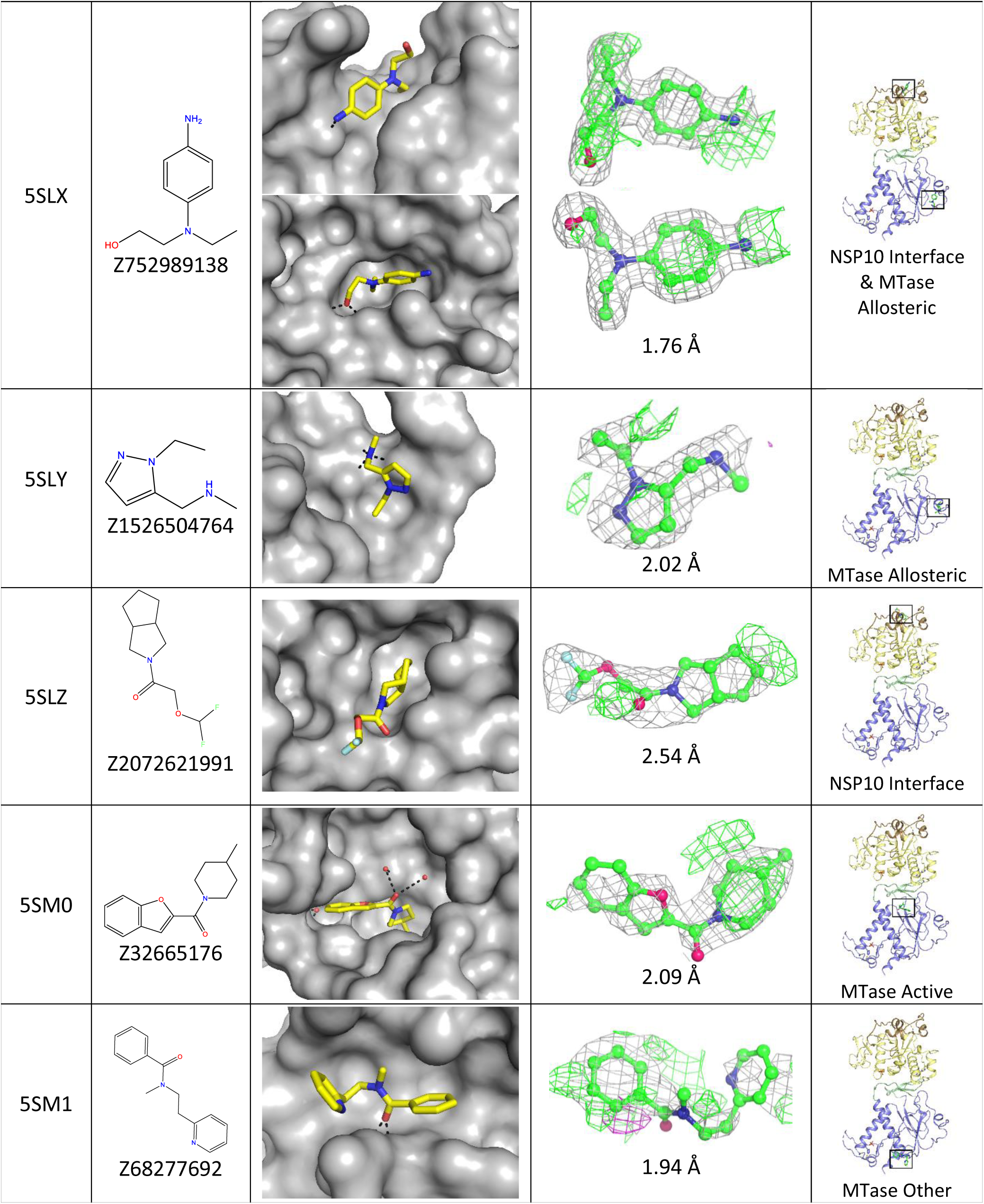
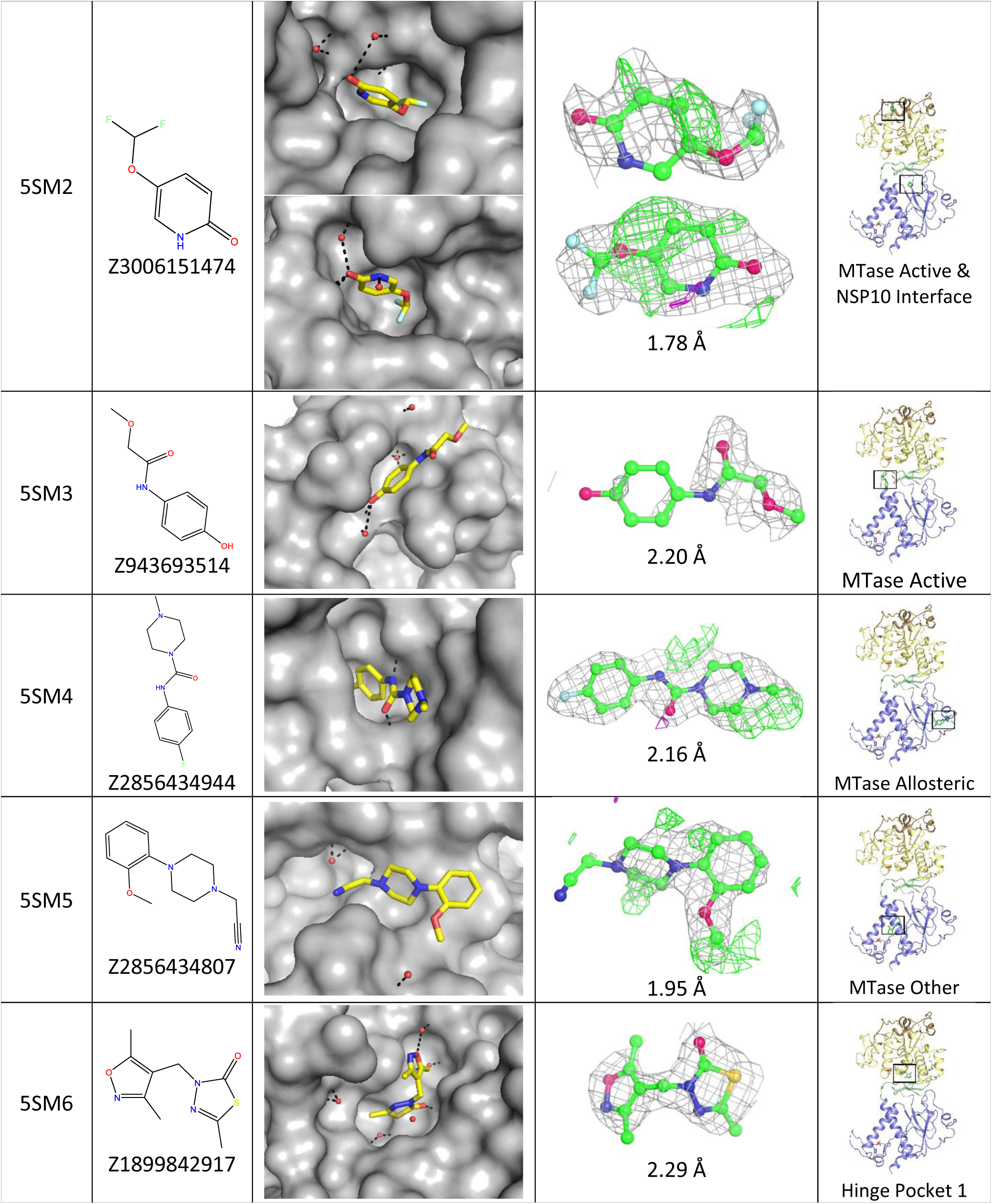
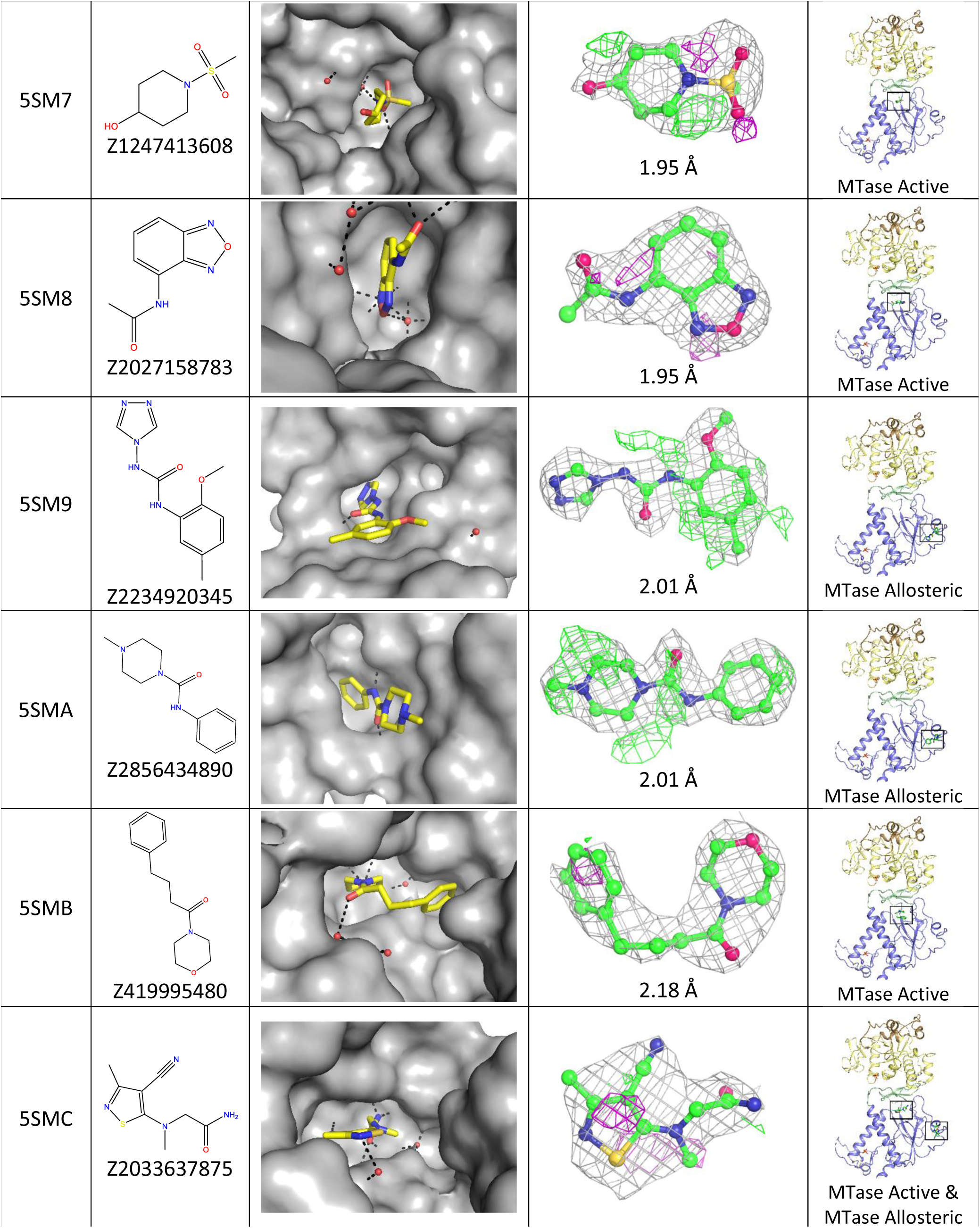
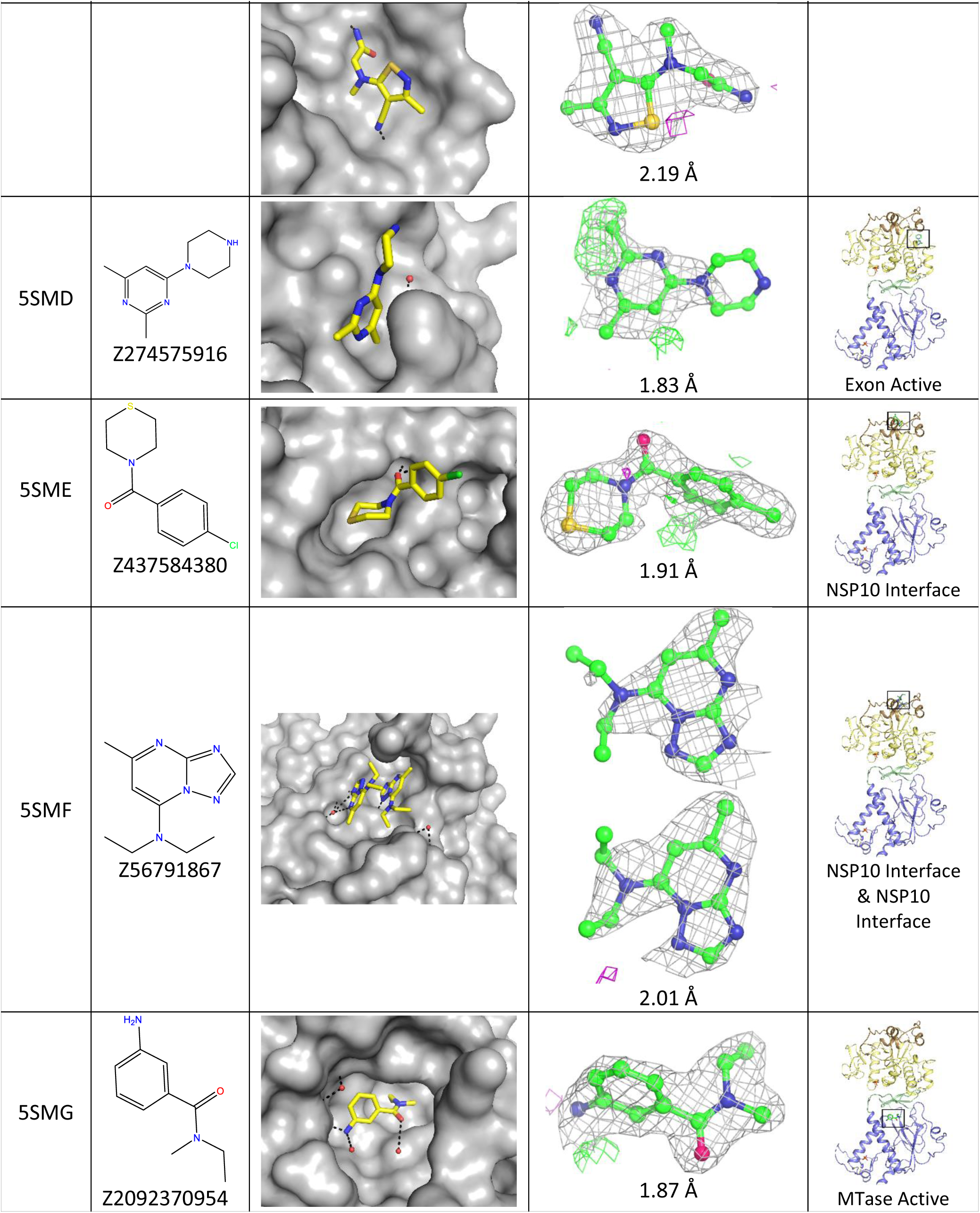
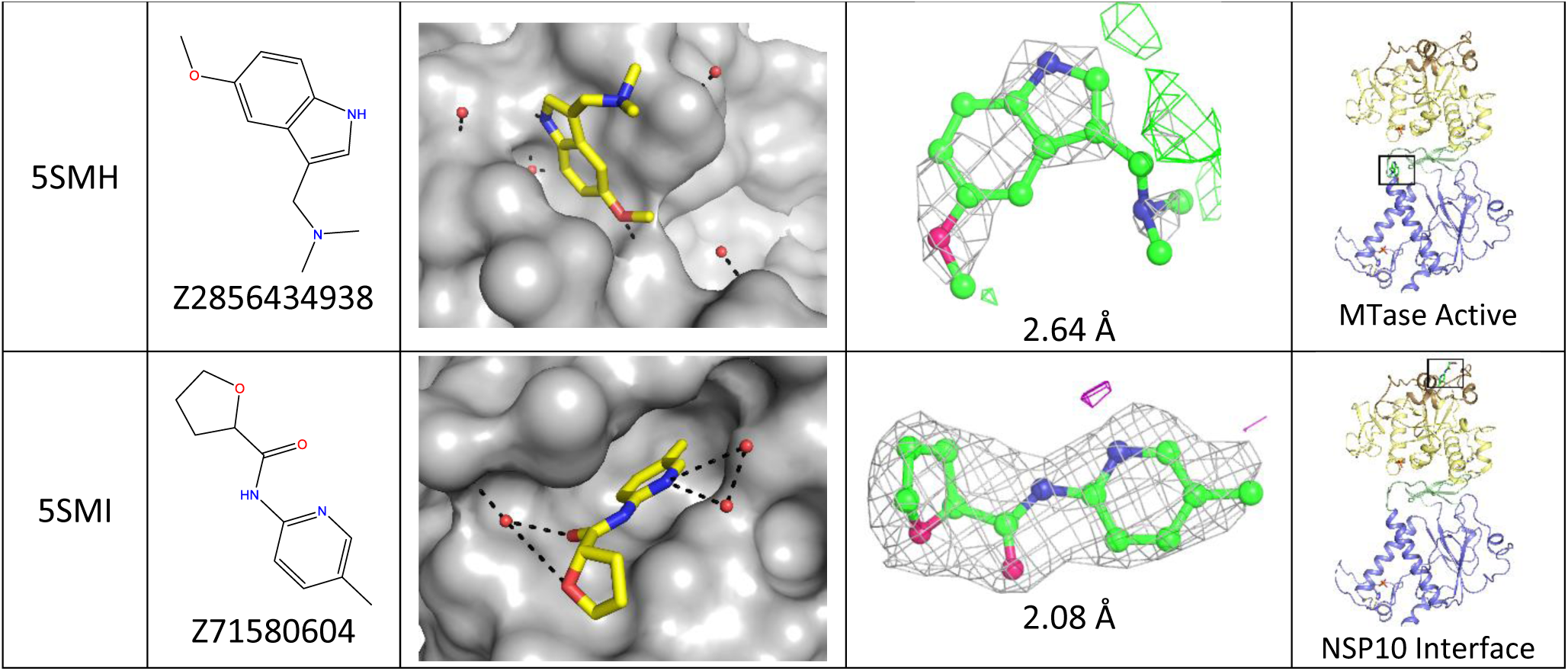
Details of all the fragment structures included in this study showing PDB codes, compound structure and codes, pocket details with polar contacts shown in black dashes, fragment locations and electron density maps. Maps shown are 2F_o_-1F_c_ at 0.7 σ in grey with F_o_-F_c_ difference maps contoured at +3 in green and -3 in purple with map resolution indicated.

## Notes

### Competing Interest Statement

The authors have declared no competing interest.

